# Direct Binding of Phosphatidylglycerol at Specific Sites Modulates Desensitization of a Pentameric Ligand-Gated Ion Channel

**DOI:** 10.1101/722512

**Authors:** Ailing Tong, John T. Petroff, Fong-Fu Hsu, Philipp A. M. Schmidpeter, Crina M. Nimigean, Liam Sharp, Grace Brannigan, Wayland W. L. Cheng

## Abstract

Pentameric ligand-gated ion channels (pLGICs) are essential determinants of synaptic transmission, and are modulated by specific lipids including anionic phospholipids. The exact modulatory effect of anionic phospholipids in pLGICs and the mechanism of this effect are not well understood. Using native mass spectrometry, coarse-grained molecular dynamics simulations and functional assays, we show that the anionic phospholipid, 1-palmitoyl-2-oleoyl-phosphatidylglycerol (POPG), preferentially binds to and stabilizes the pLGIC, Erwinia ligand-gated ion channel (ELIC), and decreases ELIC desensitization. Mutations of five arginines located in the interfacial regions of the transmembrane domain (TMD) reduce POPG binding, and a subset of these mutations increase ELIC desensitization. In contrast, the L240A mutant known to decrease ELIC desensitization, increases POPG binding. The results support a mechanism by which POPG stabilizes the open state of ELIC relative to the desensitized state by direct binding at specific sites.

## Introduction

Pentameric ligand-gated ion channels (pLGICs) are essential determinants of synaptic transmission, and the targets of many allosteric modulators including general anesthetics and anti-epileptics (1). These ion channels are embedded in a heterogeneous and dynamic lipid environment (2), and the presence of specific lipids fine-tunes the function of pLGICs and may play a role in regulating neuronal excitability and drug sensitivity (3–5). One nearly ubiquitous example is that of anionic phospholipids, which are known to modulate pentameric ligand-gated ion channels (pLGICs) such as the nicotinic acetylcholine receptor (nAchR) (6), as well as inward rectifying potassium channels (7), K(2P) channels (8), voltage-gated potassium channels (9, 10), and cyclic nucleotide-gated channels (11). In pLGICs, anionic phospholipids have been shown to shift the conformational equilibrium of the channel from an uncoupled or desensitized state to a resting state, in which agonist binding is effectively coupled to channel activation (12–14).

Studies of lipid modulation of ion channel function including modulation of pLGICs have focused on two central questions: 1) what is the exact effect of the lipid on channel function and structure, and 2) is the effect attributable to direct binding of the lipid at specific sites? *Torpedo* nAchR channel activity measured from flux assays (6, 15, 16) and agonist-induced conformational changes (13, 17) depend on anionic phospholipids. However, only a few studies have employed fast solution changes to measure current responses of pLGICs in model membranes (18), which is necessary to distinguish the effect of lipids on channel gating, specifically transitions between resting, open and desensitized states. With regard to lipid binding, early studies using electron paramagnetic resonance (EPR) of spin-labeled lipids or lipid-induced modification of fluorescent probes revealed an immobilized layer of lipids surrounding nAchRs that is enriched for certain phospholipids (19, 20) with lipids occupying specific sites (21, 22). These approaches are, however, an indirect means to examine lipid binding to ion channels. More recently, crystal structures of the pLGIC, Gloeobacter ligand-gated ion channel (GLIC), revealed bound, co-purified phospholipids in a putative open structure, and the absence of one of these phospholipids in a locally-closed structure (23, 24). Similarly, a putative desensitized structure of GLIC with a bound polyunsaturated fatty acid showed loss of the aforementioned phospholipid density that is bound to the open state (25). Both of these studies suggest that bound phospholipids at specific sites stabilize the open state of the channel, although the identity of these lipids remains unknown. Furthermore, the absence of a lipid density in a crystal structure is not necessarily an indication of lack of binding.

Native mass spectrometry (MS) has proven to be a powerful tool to directly measure binding of endogenous and exogenous lipids to membrane proteins (26, 27). In addition, coarse-grained molecular dynamics (MD) simulations provide a complementary approach to examine lipid interactions with membrane-embedded pLGICs at time scales that allow equilibration of lipid binding sites (28, 29). We sought to determine whether phospholipids bind directly and selectively to a pLGIC by native MS and coarse-grained MD simulations, and whether specific binding interactions modulate channel function by measuring ELIC activity in liposomes of defined lipid composition. Erwinia ligand-gated ion channel (ELIC), a prototypical pLGIC and biochemically tractable target, is also sensitive to its lipid environment. ELIC was found to be inactive when reconstituted in 1-palmitoyl-2-oleoyl-phosphatidylcholine (POPC) membranes fused to *Xenopus* oocyte membranes, similar to the nAchR (30). After optimizing native MS for ELIC, we demonstrate that phospholipids directly bind to ELIC, with more binding observed for the anionic phospholipid, 1-palmitoyl-2-oleoyl-phosphatidylglycerol (POPG) compared to zwitterionic phospholipids, 1-palmitoyl-2-oleoyl-phosphatidylethanolamine (POPE) and POPC. Consistent with this finding, coarse-grained simulations of ELIC in a lipid bilayer show enrichment of annular POPG compared to POPC or POPE. In addition, POPG selectively stabilizes ELIC against thermal denaturation indicative of a specific binding interaction, and reduces channel desensitization. Mutations of five arginines at the transmembrane domain (TMD) intracellular and extracellular interfaces decrease POPG binding while a subset of these mutations increase desensitization. Likewise, the L240A mutant, which reduces desensitization, increases POPG binding. The results support the hypothesis that anionic phospholipids stabilize the open state of pLGICs by direct binding to sites in the TMD adjacent to the lipid-facing transmembrane helix 4 (TM4) (3).

## Results

### Selective binding of phospholipids to ELIC

Native MS of ELIC purified in dodecyl maltoside (DDM) was optimized on a Q-Exactive EMR mass spectrometer as previously described (27). Optimal desolvation of the pentamer required activation energies that resulted in some dissociation into tetramer and monomer (Fig. 1A). Nevertheless, both the pentamer and tetramer species showed multiple bound small molecules of ∼750 Da, likely corresponding to co-purified phospholipids (up to 8 and 6 lipids per multimer were observed for the pentamer and tetramer, respectively) (Fig. 1A). To determine the identity of these lipids, we performed a lipid extraction from the purified ELIC preparation, and analyzed the sample using tandem MS. This revealed multiple PE and PG phospholipids with different acyl chains that mirror the phospholipids extracted from *E. coli* membranes (Supplementary Table 1). Quantification of the MS intensities for PG relative to PE species yielded a higher relative abundance of PG co-purified with ELIC compared to *E. coli* membranes, suggesting that ELIC preferentially binds PG in its native environment (Supplementary Fig. 1, Supplementary Table 1).

**Figure 1.**
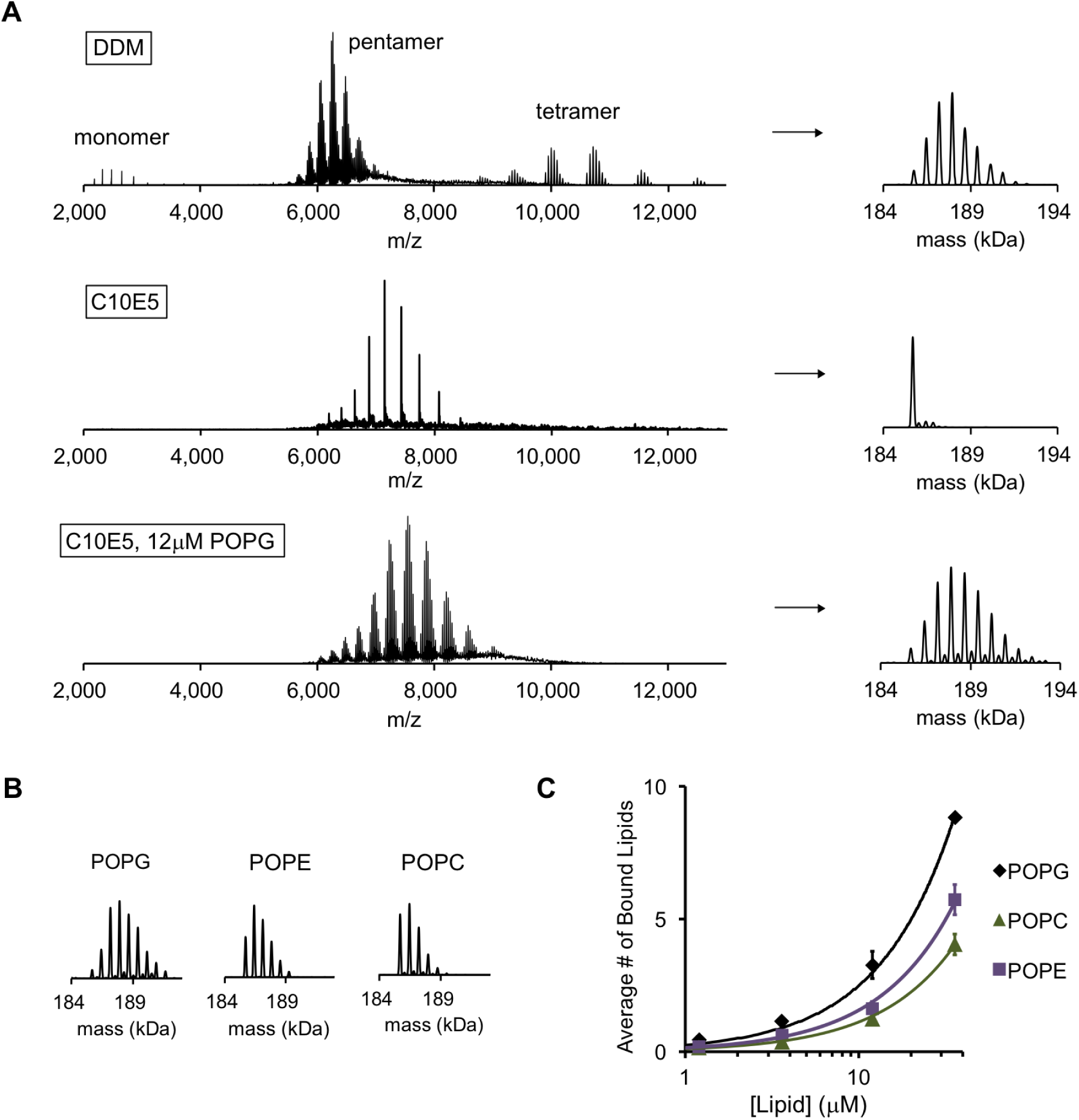
POPG binds selectively to ELIC. (**A**) Native MS spectra of ELIC in DDM, C10E5, and C10E5 with 12 μM POPG. *Left* shows full spectra and *right* shows deconvoluted spectra. (**B**) Deconvoluted spectra of ELIC in 12 μM of the indicated phospholipid. (**C**) Plot of the average number of bound phospholipids per pentamer at varying concentrations of POPG, POPE and POPC (n=3-6, ±SD).

To examine direct binding of exogenous phospholipids to ELIC, we performed a detergent screen to delipidate ELIC focusing on detergents that are also superior for native MS measurements (31). The polyethylene glycol-based detergent, C10E5, proved best for this application, yielding a stable, delipidated pentamer by native MS with lower charge states and no dissociation of the pentamer (Fig. 1A). This observation is consistent with previous reports in other membrane proteins (31, 32). Addition of varying concentrations of the anionic phospholipid, POPG, to 1 μM ELIC showed concentration dependent binding (Supplementary Fig. 2). We quantified this binding by calculating the average number of bound phospholipids at each concentration. For example, at 12 μM POPG, native MS spectra revealed up to 9 bound POPG per pentamer or an average of 2.9 POPG per pentamer (Fig. 1B and 1C). The average number of bound POPG was equivalent for most charge states, and decreased modestly at charge states higher than +26 likely due to electrostatic repulsion within the ELIC-POPG complexes (Supplementary Fig. 3); therefore, deconvolution was performed for charge states +26 and lower. Less binding was observed for the neutral phospholipids, POPE and POPC (Fig. 1B and 1C), indicating that the anionic phospholipid, POPG, either binds with higher affinity or at more sites.

To further examine phospholipid interactions with ELIC using a molecular model, we performed coarse-grained MD simulations on binary POPG/POPC and POPG/POPE model membranes containing a single ELIC pentamer (Fig. 2A). Unlike fully-atomistic simulations, coarse-grained simulations permit significant diffusion of lipids over simulation time scales. The boundary lipid composition can thus equilibrate over the simulation time, even if it varies significantly from the bulk membrane composition. The POPG fraction was varied between 0 and 70%. Enrichment or depletion of POPG among boundary lipids for each concentration was quantified using the boundary lipid metric B (Equation 7, see Methods). For a given lipid species, B>0 reflects enrichment, B<0 reflects depletion, and B=0 reflects random mixing. For POPG, B>0 for all compositions tested (Fig. 2B). This result indicates that if POPG is present in the membrane, it is enriched among boundary lipids. This enrichment is strongest for lower amounts of POPG (i.e. lower *x_PG_*), consistent with specific binding of POPG to ELIC.

**Figure 2.**
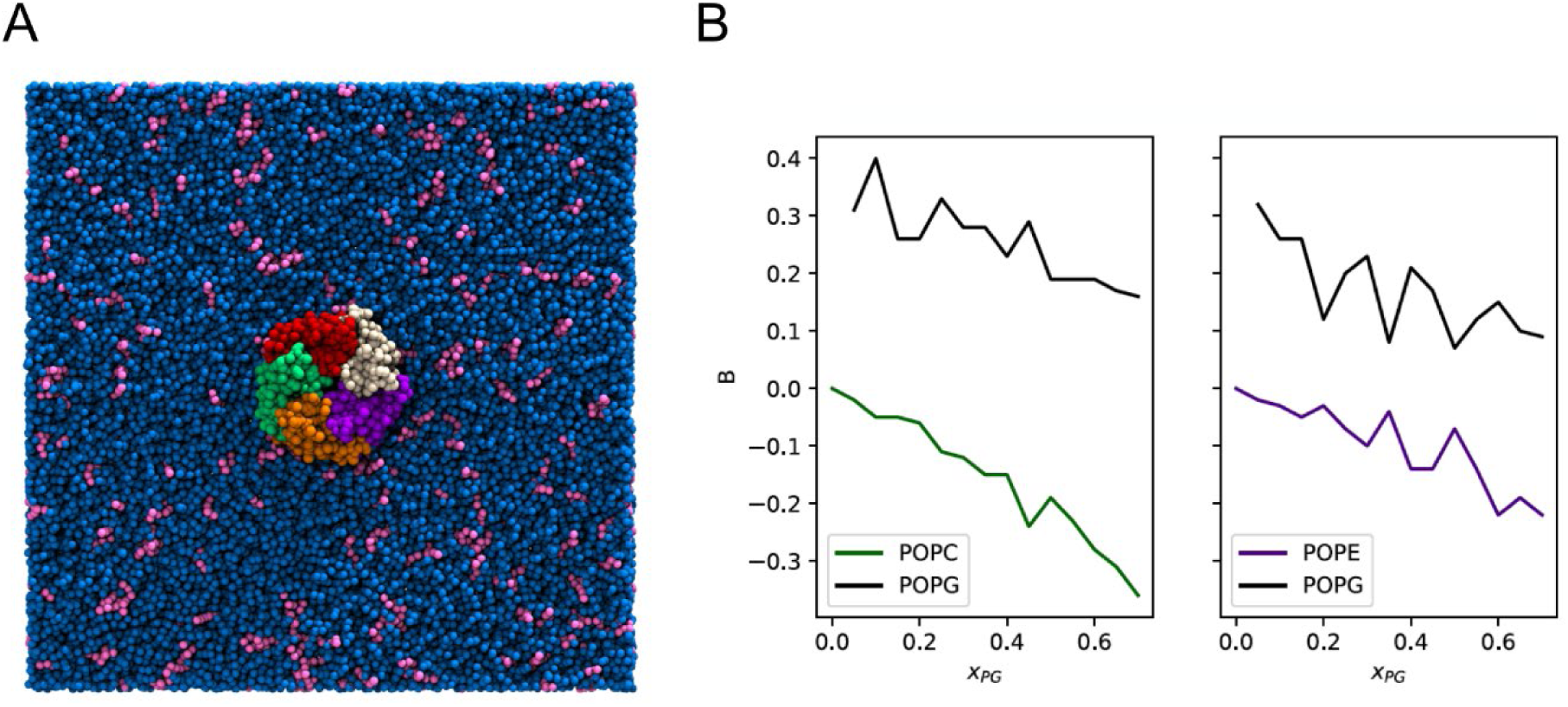
Enrichment of POPG among ELIC boundary phospholipids from coarse-grained simulations. (**A**) Image of the simulation model of ELIC embedded in a membrane consisting of 10% POPG (pink) and 90% POPC (blue). The view is from the extracellular side of ELIC perpendicular to the membrane. (**B**) The boundary enrichment metric, B, is shown for phospholipid species in POPC/POPG membranes (left) or POPE/POPG membranes (right) over a range of POPG mole fractions (*x*_*PG*_). B is defined in Equation 7 (see Methods) and reflects the fractional difference between the amount of a lipid species found in the boundary and the bulk membrane: B>0 indicates enrichment, B<0 indicates depletion, and B = 0 indicates no difference in mole fraction between the bulk and the boundary.

The average number of boundary phospholipids was 31.6 ± 2.5 (±SD) across all compositions, and the total did not vary systematically with membrane composition. Therefore, we assumed that the stoichiometries of binding for these phospholipids to ELIC are similar, and fit the native MS binding data for each phospholipid to a binomial distribution binding model with 32 binding sites of equivalent affinity (see Methods). While this is an oversimplification of phospholipid binding to ELIC in a membrane, it provides a reasonable approximation to the MS data, and reveals that POPG binds to ELIC with ∼1.9x and 2.8x higher affinity than POPE and POPC, respectively (Supplementary Fig. 4). Overall, we conclude that POPG binds to ELIC with higher affinity than POPE or POPC, resulting in POPG enrichment of annular phospholipids as seen in the coarse-grained MD simulations.

### Selective effect of POPG on ELIC stability and function

To determine the effect of POPG binding on ELIC, we first tested the stability of purified, delipidated ELIC in C10E5 against thermal denaturation in the absence and presence of POPG (33). ELIC was heated to a temperature that resulted in 85% decrease in the amplitude of the pentamer peak as assessed by size exclusion chromatography (32 °C for 15 min). POPG significantly increased the thermal stability of 1 μM ELIC with an EC_50_ (concentration of POPG for 50% effect) of 52 μM (Fig. 3A). The thermal stabilizing effect of a phospholipid was defined as the ratio of the pentamer peak height after heating with lipid versus no lipid. In contrast, POPE and POPC had no effect on ELIC stability (Fig. 3A), indicating that POPG binding selectively stabilizes the structure of ELIC. Having performed our POPG binding experiment and thermal stability assay under the same conditions, it is possible to relate the average number of bound POPG to its stabilizing effect. 36 μM POPG was the highest concentration for which the average number of bound POPG could be determined due to the overlapping of charge states from lipid-bound species (Fig. 1A, Supplementary Fig. 2). Although POPG binding does not approach saturation at this concentration, extrapolation of POPG binding and relating this extrapolation to the thermal stabilizing effect provides an approximation of the number of bound POPG needed to stabilize ELIC against thermal denaturation. Supplementary Figure 5 shows a relationship between the number of bound POPG and the stabilizing effect, which was derived by equating the POPG concentration from the functions of POPG binding (Fig. 1C) and thermal stability data (Fig. 3A). The relationship estimates that 32 POPG (number of annular lipids in ELIC from MD simulations) yields ∼82% of the thermal stabilizing effect (Supplementary Fig. 5).

**Figure 3.**
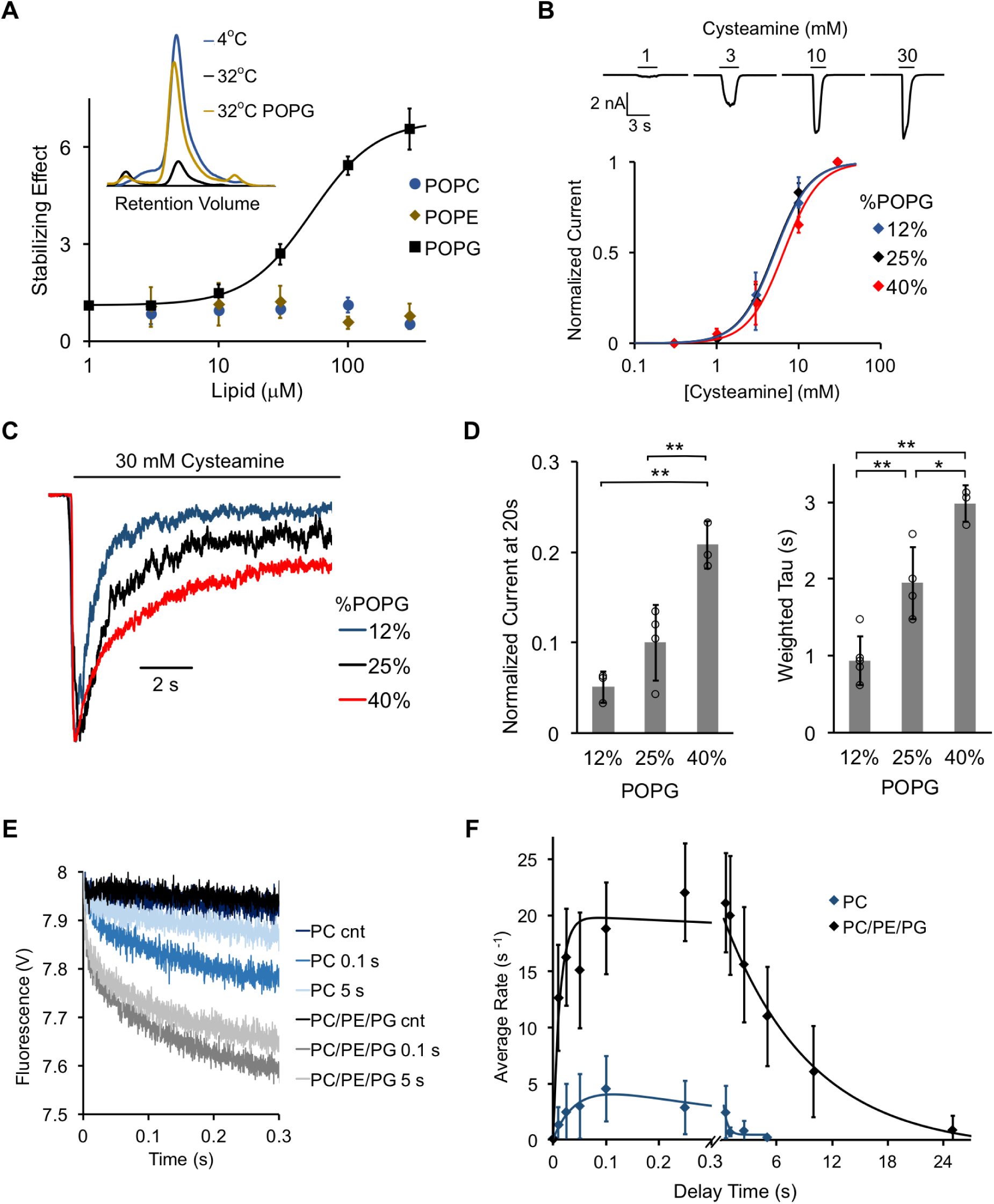
POPG selectively thermally stabilizes ELIC and decreases ELIC desensitization. (**A**) Plot of stabilizing effect (defined as the ELIC pentamer peak height with phospholipid relative to control after heating) versus phospholipid concentration (n=3, ±SD; EC_50_ = 52 μM, Hill n = 1.7). Inset shows size exclusion chromatography (SEC) profile in absorbance units of the ELIC pentamer treated at 4°C, 32°C, and 32°C with 100 μM POPG. (**B**) *Top:* Representative ELIC current responses to 30 mM cysteamine in 25 mole% POPG liposomes. *Bottom:* Normalized plots of peak current responses of ELIC to cysteamine in giant liposomes with varying mole% POPG (n=3-5, ±SD). Data are fit to Hill equation with n=2. (**C**) Representative ELIC currents in response to 30 mM cysteamine in liposomes with varying mole% POPG. (**D**) *Left:* ELIC currents 20 s after application of 30 mM cysteamine normalized to peak response at varying mole% POPG (n=4-5, ±SD, **p<0.01). *Right:* Weighted tau (time constants) of ELIC desensitization at varying mole% POPG (n=3-5, ±SD, **p<0.01, *p<0.05). (**E**) Representative fluorescence-quench time courses from sequential mixing stopped-flow experiments of ELIC in POPC liposomes or 2:1:1 POPC:POPE:POPG liposomes. Proteoliposomes were mixed with no cysteamine (cnt) or 5 mM cysteamine with a 0.1 or 5 s delay prior to mixing with Tl^+^. Note that the control traces are superimposed. (**F**) Rate constants extracted from quench kinetics as shown in (E) as a function of the incubation time with cysteamine. Data are fit with a double exponential yielding activation and desensitization time constants (n=3, ±SD).

Next, we assessed the effect of POPG on ELIC function by reconstituting the channel in giant liposomes. Optimal formation of giant liposomes was achieved using a 2:1:1 ratio of POPC:POPE:POPG (i.e. 25 mole% POPG). In this lipid membrane composition, robust ELIC currents were elicited with excised patch-clamp recordings using the agonist, cysteamine, with a peak dose response EC_50_ of 5.1 mM (Fig. 3B, Table 1, Supplementary Fig. 6A). Patch-clamp recordings were performed with 0.5 mM BaCl_2_ in the pipette and bath, which is predicted to result in an increase in the EC_50_ of cysteamine response (34). Near saturating currents were achieved at 30 mM cysteamine at which ELIC activated and desensitized with time constants of 134 ms and 1.9 s, respectively (Fig. 3C and 3D, Table 1, Supplementary Fig. 6B). These values are comparable to previous reports of outside-out patch-clamp recordings in HEK cells or oocytes (35, 36). ELIC desensitization showed complex kinetics where the majority of recordings were best fit with a double exponential and some by a single exponential. To combine data from all traces, weighted average time constants from double exponential fits were averaged with time constants from single exponential fits. The extent of desensitization was examined by measuring currents after 20 s of cysteamine application. To examine the effect of POPG on ELIC gating, excised patch-clamp recordings were performed in liposomes containing 12%, 25%, and 40% POPG. Increasing the mole% of POPG had no significant effect on cysteamine EC_50_ values or activation kinetics (Fig. 3B, Table 1, Supplementary Fig. 6), but reduced the rate and extent of desensitization (Fig. 3C and 3D, Table 1).

**Table 1:** ELIC WT channel properties in giant liposomes composed of varying mole% POPG (n=3-5, ±SD). The rate and extent of desensitization are reported as weighted time constants (*τ*), and the current after 20 s of 30 mM cysteamine application normalized to peak response. Also shown are activation time constants (*τ*) in response to 30 mM cysteamine and EC_50_s for cysteamine activation.

|  | Desensitization |  | Activation | Dose Response |
| --- | --- | --- | --- | --- |
| POPG | Weighted $\tau$ (s) | Norm Current at 20s | $\tau$ (ms) | EC<sub>50</sub> (mM cysteamine) |
| 12% | 0.93 ± 0.32 | 0.05 ± 0.02 | 112 ± 29 | 5.3 ± 1.0 |
| 25% | 1.95 ± 0.47 | 0.10 ± 0.04 | 134 ± 50 | 5.1 ± 1.2 |
| 40% | 2.98 ± 0.24 | 0.21 ± 0.03 | 133 ± 66 | 6.5 ± 1.3 |

To examine ELIC activity in the absence of POPG, a fluorescence-based stopped-flow flux assay was performed (37). ELIC was reconstituted into either POPC-alone or 2:1:1 POPC:POPE:POPG liposomes encapsulating the fluorophore ANTS (8-Aminonaphthalene-1,3,6-Trisulfonic acid). In a first mixing step, liposomes were incubated with 5 mM cysteamine to activate the channel for different amounts of time (10 ms to 25 s) after which a second mixing step was performed with Tl^+^ containing buffer. Tl^+^ can permeate through activated channels into the liposomes where it quenches ANTS fluorescence. The quenching kinetics are a measure of the channel activity upon cysteamine exposure for defined incubation times (Fig. 3E). In POPC liposomes, ELIC showed less cysteamine-elicited ion flux compared to ELIC in POPC:POPE:POPG liposomes (Fig. 3E and 3F, Table 2), as estimated from the overall rate of Tl^+^ flux. The rate of activation was modestly faster in POPC:POPE:POPG liposomes compared to POPC (Fig. 3F, Table 2). More strikingly, the rate of desensitization was more than 20-fold faster in POPC liposomes, leading to a decrease in the lifetime of the open state (Fig. 3E and 3F, Table 2).

**Table 2:** ELIC WT activation and desensitization rate constants derived from a double exponential fit to the time course of flux in Fig. 3F (n=3, +/- SD).

|  | Activation Rate Constant | Desensitization Rate Constant |
| --- | --- | --- |
| POPC | 24 ± 8 s<sup>-1</sup> | 2.4 ± 0.08 s<sup>-1</sup> |
| POPC:POPE:POPG (2:1:1) | 75 ± 22 s<sup>-1</sup> | 0.11 ± 0.04 s<sup>-1</sup> |

In summary, POPG selectively increases the thermal stability of ELIC, and modulates channel activity by stabilizing the open relative to the desensitized state. We hypothesize that POPG decreases receptor desensitization by direct binding at specific sites.

### Five interfacial arginines contribute to POPG binding

In other ion channels, guanidine groups from interfacial arginine side chains are thought to mediate binding of anionic phospholipids by charge interactions with the phospholipid headgroup (9, 38). To test the hypothesis that this mechanism is present in a pLGIC, we mutated all five arginines in the inner and outer interfacial regions of the ELIC TMD to glutamine (Fig. 4A). Phospholipid binding was then assessed by delipidating each mutant in C10E5, and measuring binding of POPG by native MS. While R123Q, R286Q, and R299Q could be stably delipidated, R117Q and R301Q aggregated (Fig. 4B). However, we found that double mutants with R299Q (i.e. R117Q/R299Q and R301Q/R299Q) could be stably delipidated. Thus, double mutants of all arginine mutants in combination with R299Q were expressed and delipidated (Fig. 4B). In the presence of 12 μM POPG, the single mutants showed moderate decreases (13-18%) in the average number of bound POPG compared to WT (Fig. 4B). This decrease was not statistically significant. However, all double mutants significantly decreased the average number of bound POPG relative to WT by 38-41% and relative to R299Q by 27-32% (Fig. 4B). These results indicate that each interfacial arginine contributes approximately equally to POPG binding in ELIC. It is likely that significant decreases in binding could only be appreciated in the double mutants because of the variability in the data.

**Figure 4.**
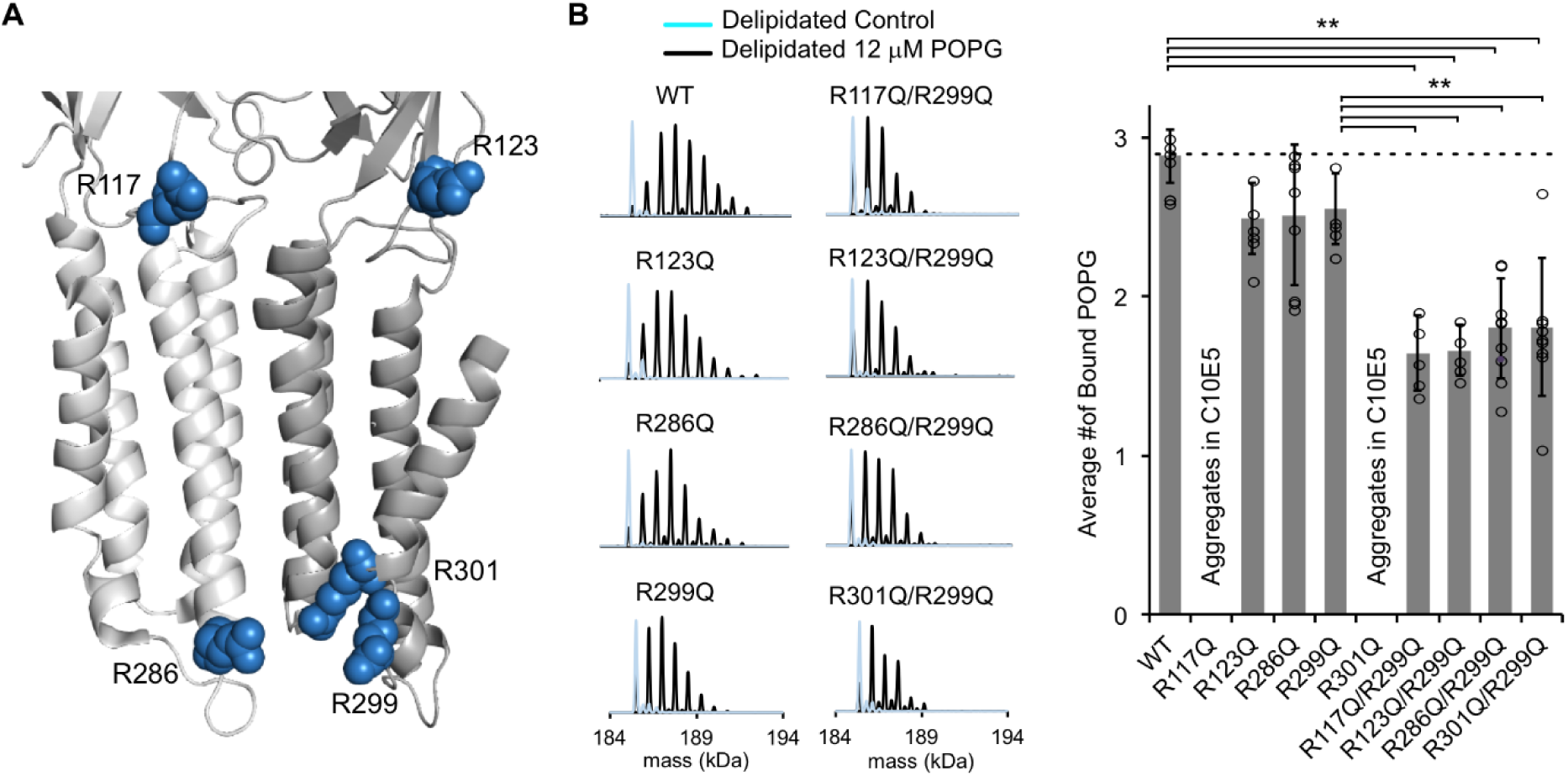
Mutations of five interfacial arginines reduce POPG binding. (**A**) Structure of ELIC showing two adjacent subunits and five arginine side chains that were mutated to glutamine. (**B**) *Left:* Representative deconvoluted spectra of ELIC WT and indicated mutants. Blue indicates spectra of delipidated ELIC in C10E5. Black indicates spectra of delipidated ELIC in C10E5 with 12 μM POPG. *Right:* Plot of average number of bound POPG for ELIC WT and mutants, delipidated in C10E5, with 12 μM POPG (n=5-8, ±SD, **p<0.01).

We further examined these sites of interaction using our coarse-grained MD simulations. To identify whether boundary POPG were localized around specific helices or residues, two-dimensional densities of the negatively-charged headgroup bead were calculated. The distributions are separated by leaflet where each leaflet contained 10% POPG. As shown in Figure 5A, POPG was more likely to interact with ELIC in the inner leaflet than the outer leaflet, consistent with three out of five interfacial arginine residues being located on the intracellular interface of the ELIC TMD. These three arginines are located on TM3 (R286) and TM4 (R299 and R301). Contacts between POPG and all three of these residues are also visible in individual frames of the simulation (Fig. 5B). Moreover, POPG is more likely to be contacting the interfacial residues in TM4 (such as R299 and R301) than accessible interfacial residues in any other helix (Fig. 5A). The remaining two arginine residues are located at the TMD-ECD interface (R117 and R123). POPG density in the outer leaflet localized to these residues at intrasubunit sites between TM4 and TM1 or TM4 and TM3 (Fig. 5A), and contacts between these residues and POPG headgroups in the outer leaflet were also observed in snapshots from the MD simulations (Fig. 5B). In summary, the native MS data and coarse-grained MD simulations demonstrate that five interfacial arginines contribute to specific POPG binding sites in the inner and outer leaflets adjacent to TM4.

**Figure 5.**
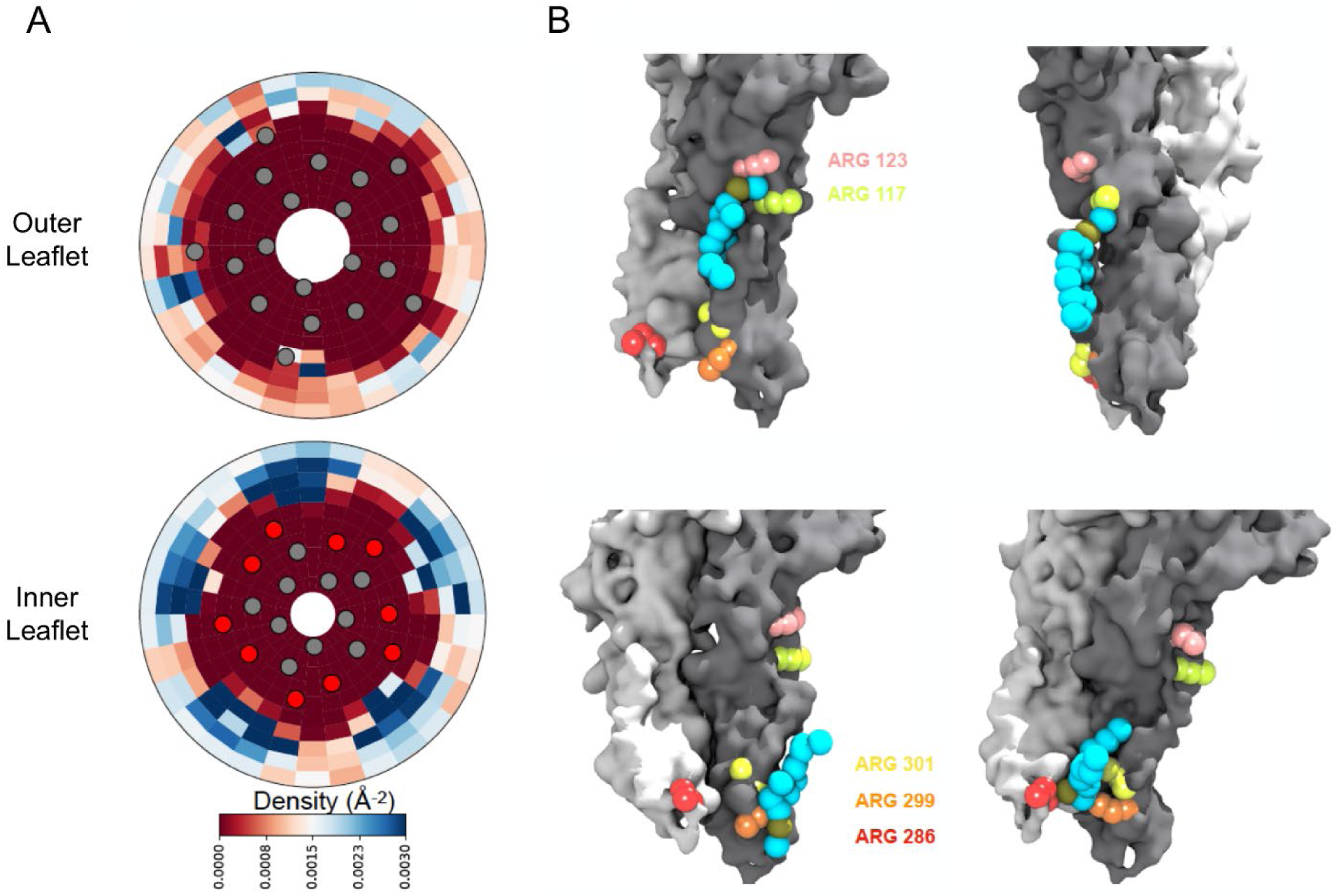
Density calculations of lipids in binary membranes and visualization of direct POPG-ELIC interactions at 10% POPG. (**A**) Distribution of POPG density in a POPG-POPC membrane, within 40 Å from the ELIC pore over the last half of a 15 µs simulation, for both the outer leaflet (*top*) and the inner leaflet (*bottom*). Density is colored according to the color bar, where red and blue represent low and high POPG density, respectively. Circles represent the ELIC transmembrane backbone center of mass, with the helices containing the interfacial arginines colored in red (**B**) Representative frames after ∼9 µs of simulation, showing multiple POPG binding modes associated with high density areas in (A). Two adjacent subunits of ELIC are shown in grey and white, while arginine side chains of interest are colored in peach, lime-yellow, orange, yellow, and red. POPG phosphate is colored in tan with the rest of the lipid in cyan.

### Specific interfacial arginines mediate POPG effect

Having established that ELIC selectively binds POPG over neutral phospholipids, and that binding is mediated by five interfacial arginines, we examined the role of these binding sites on ELIC desensitization. We reconstituted each single mutant into giant liposomes composed of a 2:1:1 ratio of POPC:POPE:POPG (25% POPG) to test channel function by excised patch-clamping. We hypothesized that since increasing mole% POPG decreases ELIC desensitization, certain arginine mutants, which disrupt POPG binding, may increase ELIC desensitization. Indeed, all five single arginine mutants showed variable increases in the rate or extent of desensitization; however, these differences were generally small and statistically insignificant except for R301Q (Fig. 6, Table 3). We also tested the double mutants, which showed significant decreases in POPG binding. Three double mutants (R117Q/R299Q, R123Q/R299Q, R301Q/R299Q) showed a significant increase in the extent of desensitization while two (R117Q/R299Q, R301Q/R299Q) also showed a significant increase in the rate of desensitization (Fig. 6, Table 3). The effects observed in the double mutants approximate the sum of effects observed in the single mutants. Only R286Q/R299Q did not affect the extent or rate of desensitization (Fig. 6, Table 3). The EC_50_ of cysteamine response and activation kinetics were also measured for all mutants; only R117Q and R117Q/R299Q showed significantly lower EC_50_ and activation tau values compared to WT (Table 3, Supplementary Fig. 7). Overall, these data indicate that four of five interfacial arginine residues that reduce POPG binding (i.e. R117, R123, R299, R301) also increase the rate and/or extent of ELIC desensitization.

**Figure 6.**
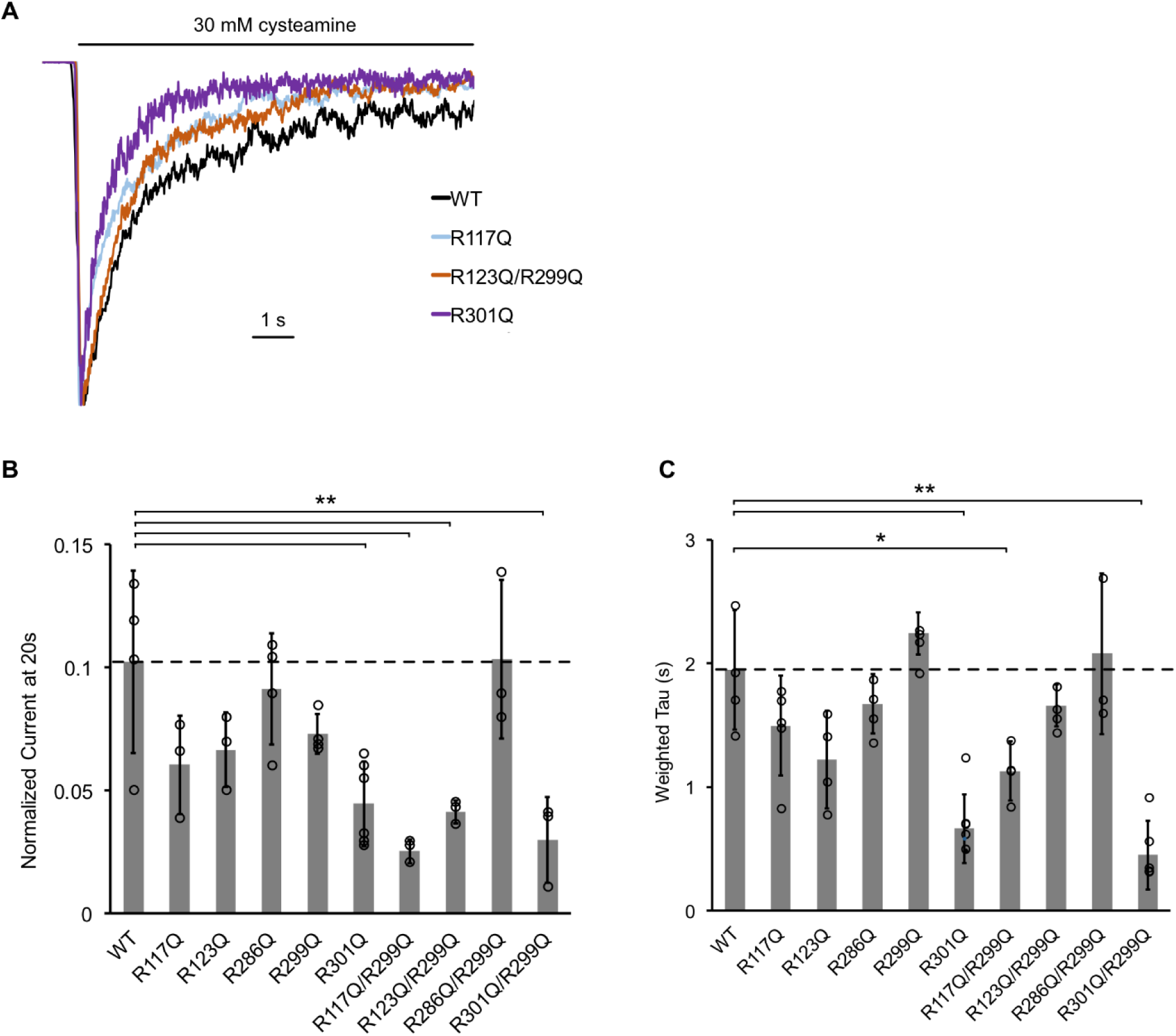
The effect of ELIC mutants on desensitization. (**A**) Normalized ELIC WT and mutant current responses to 30 mM cysteamine in 25% POPG liposomes. (**B**) Graph of ELIC WT and mutant currents 20 s after application of 30 mM cysteamine normalized to peak response in 25 mole% POPG liposomes (n=3-7, ±SD, **p<0.01, *p<0.05). (**C**) Same as (B) for weighted tau (time constants) of desensitization.

**Table 3:**
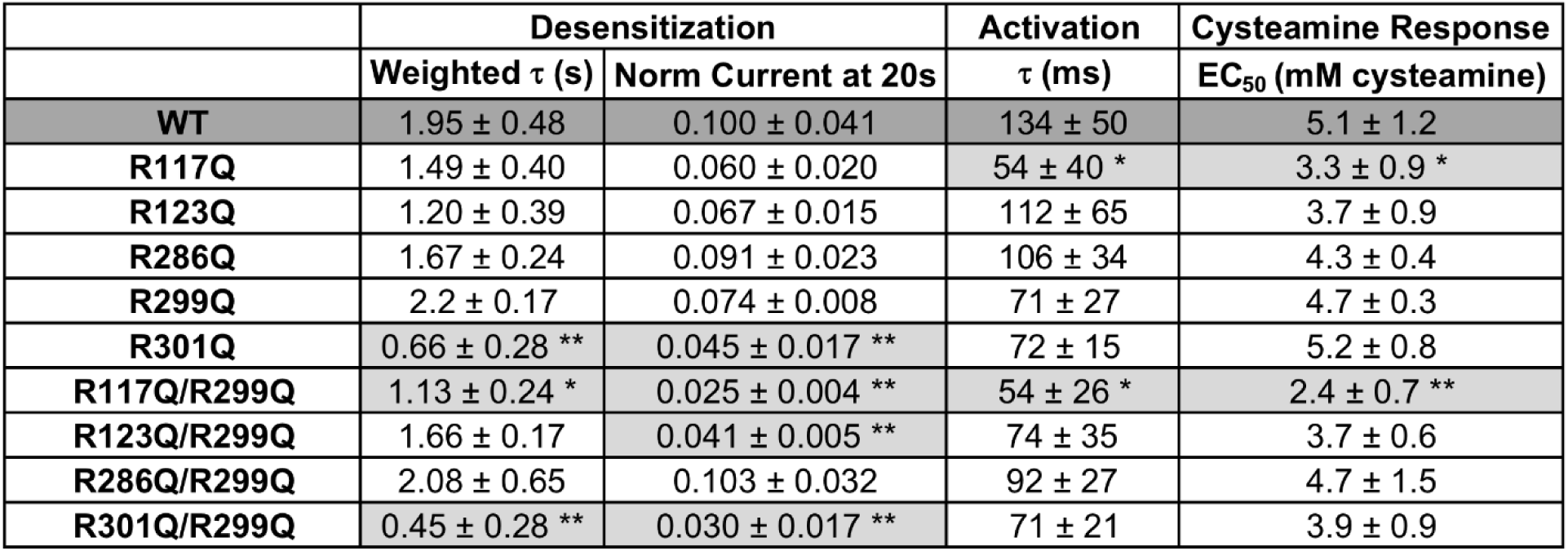
ELIC WT and mutant channel properties in giant liposomes composed of 25 mole% POPG (n=3-7, ±SD). Shown are weighted time constants (*τ*) for desensitization and currents 20 s after application of 30 mM cysteamine normalized to peak response. Also shown are activation time constants and EC_50_ of cysteamine response. Light gray indicates mutant values which are significantly different from WT (dark gray) (** p<0.01, * p<0.05).

|  | Desensitization |  | Activation | Cysteamine Response |
| --- | --- | --- | --- | --- |
| | Weighted $\tau$ (s) | Norm Current at 20s | $\tau$ (ms) | EC<sub>50</sub> (mM cysteamine) |
| WT | 1.95 ± 0.48 | 0.100 ± 0.041 | 134 ± 50 | 5.1 ± 1.2 |
| R117Q | 1.49 ± 0.40 | 0.060 ± 0.020 | 54 ± 40 * | 3.3 ± 0.9 * |
| R123Q | 1.20 ± 0.39 | 0.067 ± 0.015 | 112 ± 65 | 3.7 ± 0.9 |
| R286Q | 1.67 ± 0.24 | 0.091 ± 0.023 | 106 ± 34 | 4.3 ± 0.4 |
| R299Q | 2.2 ± 0.17 | 0.074 ± 0.008 | 71 ± 27 | 4.7 ± 0.3 |
| R301Q | 0.66 ± 0.28 ** | 0.045 ± 0.017 ** | 72 ± 15 | 5.2 ± 0.8 |
| R117Q/R299Q | 1.13 ± 0.24 * | 0.025 ± 0.004 ** | 54 ± 26 * | 2.4 ± 0.7 ** |
| R123Q/R299Q | 1.66 ± 0.17 | 0.041 ± 0.005 ** | 74 ± 35 | 3.7 ± 0.6 |
| R286Q/R299Q | 2.08 ± 0.65 | 0.103 ± 0.032 | 92 ± 27 | 4.7 ± 1.5 |
| R301Q/R299Q | 0.45 ± 0.28 ** | 0.030 ± 0.017 ** | 71 ± 21 | 3.9 ± 0.9 |

### L240A reduces desensitization and enhances POPG binding

If mutations that disrupt POPG binding increase receptor desensitization, then a mutation that decreases desensitization may enhance POPG binding. Mutation of a conserved pore-facing 9’ TM2 leucine residue is known to slow desensitization in ELIC (L240A) (36) and other pLGICs. To confirm this finding in our system, we reconstituted ELIC L240A into giant liposomes for patch-clamping and observed a significant reduction in the extent and rate of desensitization (Fig. 7A). To examine POPG binding, L240A was then de-lipidated in C10E5 for native MS. Interestingly, L240A significantly increased POPG binding compared to WT at 12 μM POPG (∼1.7x increase in average number of POPG bound) (Fig. 7B).

**Figure 7.**
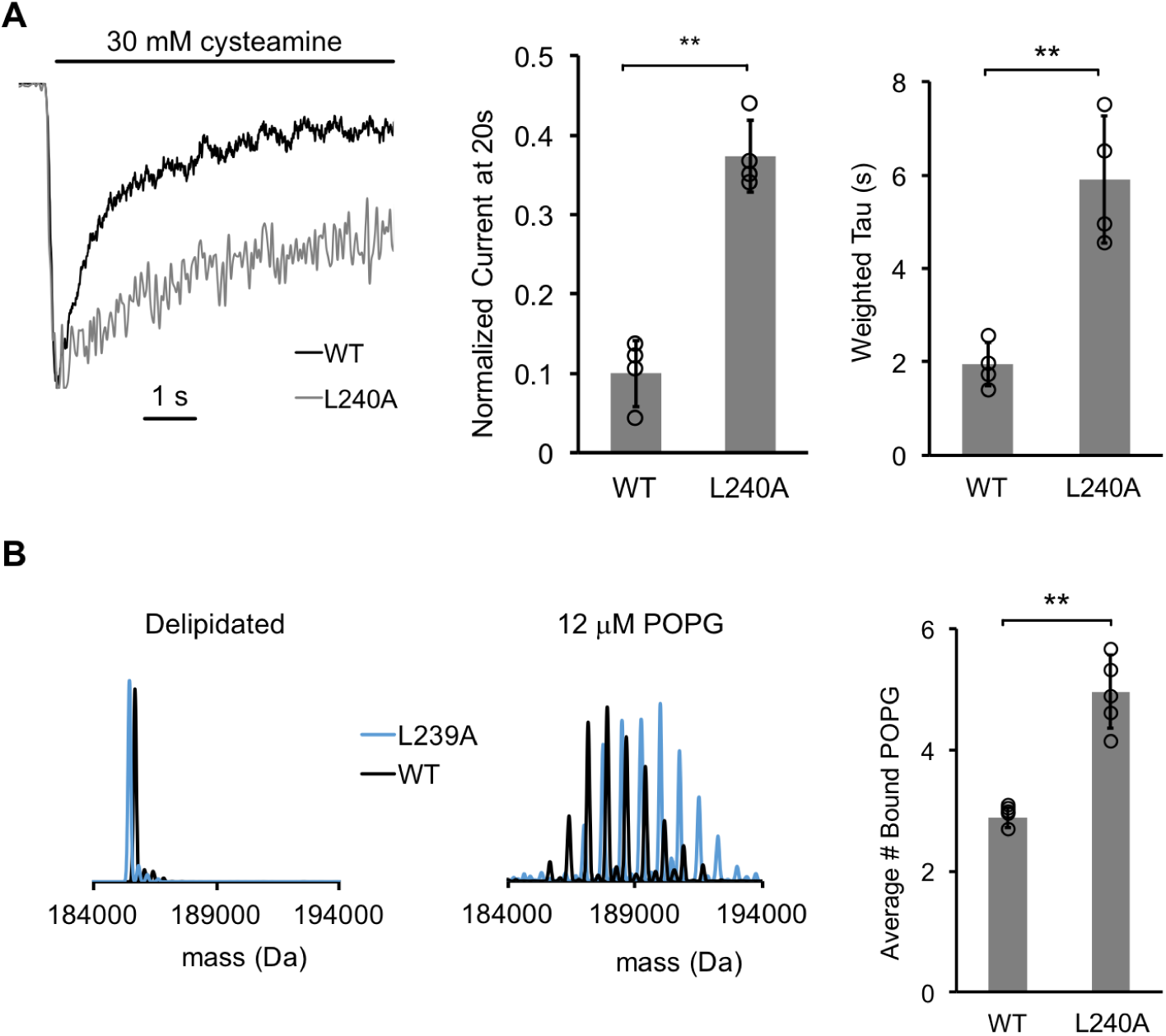
The L240A mutant decreases desensitization and increases POPG binding. (**A**) *Left:* Normalized ELIC WT and L240A current responses to 30 mM cysteamine in 25% POPG liposomes. *Middle:* ELIC WT and L240A currents 20 s after application of 30 mM cysteamine normalized to peak response at 25 mole% POPG (n=4-5, ±SD, **p<0.01). *Right:* Weighted tau (time constants) of ELIC WT and L240A desensitization time courses at 25 mole% POPG (n=4-5, ±SD, **p<0.01). (**B**) *Left:* Representative deconvoluted spectra of ELIC WT (black) and L240A (blue) showing ELIC delipidated in C10E5 without and with 12 μM POPG. *Right:* Graph of average number of bound POPG for ELIC WT and L240A, delipidated in C10E5, with 12 μM POPG (n=4-5, ±SD, **p<0.01).

## Discussion

Recent structural and computational evidence suggests that lipids bind to pLGICs at specific sites within the TMD (24, 25, 39–41). However, there is a scarcity of evidence showing that changes in direct lipid binding are correlated with functional effects (25). We show that the anionic phospholipid, POPG, selectively binds to ELIC using native MS, thermally stabilizes the channel, and decreases receptor desensitization. Overall, these data support the idea that lipid binding directly affects receptor stability and function. Further, mutations of arginine residues that reduce POPG binding also increase ELIC desensitization to varying degrees. While it is possible that these arginine mutations increase desensitization through a mechanism other than their effect on POPG binding, the correlation between binding and desensitization, and the finding that the L240A mutation, which reduces desensitization, increases POPG binding affinity supports this conclusion. Remarkably, the L240A mutation, which is located in the channel pore and remote from the lipid interface (Fig. 8), appears to allosterically alter the affinity of ELIC for POPG. Lipids may modulate ion channel activity through indirect effects on the physical properties of the membrane or through direct binding interactions (42, 43). The lipid binding data presented in this study using native MS provides evidence that direct binding of anionic phospholipids allosterically stabilizes the open state of a pLGIC relative to the desensitized state.

**Figure 8.**
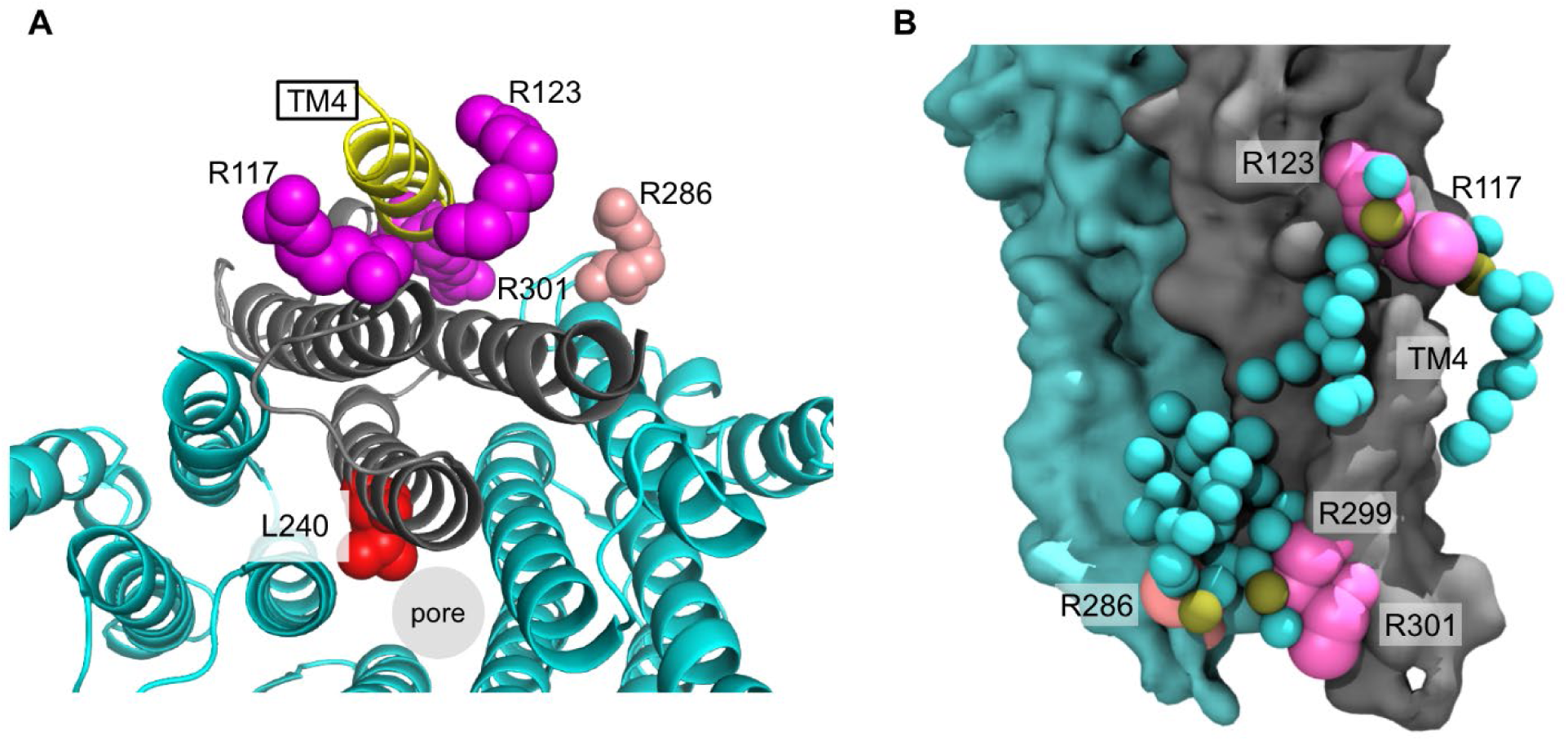
Arginines involved in POPG binding and ELIC desensitization. **(A)** Top view of ELIC highlighting TM4 (yellow) and showing the side chains of R117, R123 and R301 (magenta) adjacent to TM4, which increase ELIC desensitization, and R286 at the subunit interface (salmon), which has no effect on desensitization. L240 (red) faces the pore. **(B)** Image from coarse-grained simulations with 50% POPG showing two adjacent ELIC subunits and the mutated arginine side chains (R117, R123, R299 and R301 in magenta; R286 in salmon). Also, shown are all POPG lipids making contacts with the TMD in this snapshot.

Membrane proteins including pLGICs are thought to determine their lipid microenvironment by specific binding interactions (28, 44). Our native MS measurements provide unique insights into phospholipid interaction with a pLGIC. First, we find that more POPG binds to ELIC compared to POPE or POPC at equivalent concentrations, suggesting that POPG binds to ELIC with higher affinity. This is supported by enrichment of POPG compared to POPE in phospholipids that are co-purified with ELIC, and coarse-grained simulations which show enrichment of POPG among the boundary phospholipids of ELIC. Second, native MS also allows determination of the stoichiometry and sites of lipid binding (32, 45). By relating binding stoichiometry and thermal stability, the data estimate that 32 POPG lipids, which is the average number of annular lipids in ELIC from MD simulations, results in greater than 80% of the stabilizing effect against thermal denaturation (Supplementary Fig. 5). This suggests that maximal thermal stability is achieved when the entire ELIC transmembrane domain (TMD) is surrounded by POPG. Although five interfacial arginine residues were identified to contribute to POPG binding in ELIC (25 arginines total), it is conceivable that each arginine side chain may interact with more than one phospholipid headgroup or that other sites exist.

To quantify the effect of the ELIC double mutants on specific POPG binding sites, we also fit the native MS binding data for the double mutants to a binomial binding model using the dissociation constant for POPG binding to WT and varying the number of available sites. POPG binding to the double mutants was best fit with a reduction in the number of available sites from 32 in WT to 18-21 in the mutants (Supplementary Fig. 4). Given the ∼35-45% decrease in bound phospholipid with each double mutant (mutation of two out of five arginines), we conclude that phospholipid binding at these residues constitute the highest affinity sites. This is indeed consistent with the POPG densities from the coarse-grained simulations, which show discrete enrichment of POPG lipids adjacent to these residues. Disruption of binding sites by mutation of these arginines may not prevent the occupancy of lipids at these sites per se, but may alter the lipid binding modes or occupancy times at these sites.

Previous studies examining the effects of lipids on pLGIC function found that nAchR and ELIC are inactive in POPC-only membranes (6, 15, 30), and it was proposed that this is due to uncoupling of agonist binding to channel activation (46). To examine ELIC channel activity in liposomes lacking anionic phospholipid, we utilized a stopped-flow flux assay, and demonstrated cysteamine-elicited flux by ELIC in POPC-only liposomes. The high sensitivity of this assay may be the reason ELIC activity could be detected, contrary to a prior study in which ELIC in POPC liposomes were injected into *Xenopus* oocytes (30). However, the ELIC activity was significantly decreased compared to POPC:POPE:POPG (2:1:1) liposomes. The low protein concentration used in this assay does not allow us to assess the reconstitution efficiencies. Thus, the overall lower flux rates and smaller amplitudes in POPC could stem from lower protein reconstitution. However, the faster desensitization kinetics in POPC liposomes can be resolved reliably, and are consistent with the patch-clamp measurements. The results substantiate the role of POPG in stabilizing the open state relative to the desensitized state, and demonstrate the utility of measuring pLGIC activity in liposomes of defined lipid composition using complementary patch-clamp and stopped-flow flux techniques.

While pLGICs are known to be sensitive to their lipid environment, the binding sites that mediate lipid modulation are not well defined. It has been proposed that TM4 is a lipid sensing structure in pLGICs due to its proximity to the lipid membrane and sensitivity to mutagenesis (41, 47–49). Furthermore, crystal structures of GLIC show bound lipids within intrasubunit grooves between TM4-TM1 and TM4-TM3 (23), which have been proposed to be important determinants of channel opening (24, 25). Photolabeling studies have also identified intrasubunit neurosteroid binding sites adjacent to TM4 that mediate neurosteroid modulatory effects (50, 51). We show that POPG binding at multiple interfacial arginine residues, including R117, R124, R299 and R301 which are localized to the extracellular and intracellular sides of TM4 (Fig. 8), are likely important in mediating the effect of POPG on ELIC desensitization. Examination of boundary POPG from coarse-grained simulations with high POPG mole% (50%) at 15 μs shows POPG headgroups making contacts with all of these arginine side chains, and illustrates potential binding modes for the acyl chains (Fig. 8B). For example, boundary POPG with headgroups that interact with R301 or R299 have acyl chains that make contacts with intrasubunit sites along the intracellular side of TM4 (Fig. 8B and 5B). R301, which has the largest effect on desensitization when mutated, is conserved among many mammalian pLGICs including GABA_A_R and nAchR isoforms, and R299 is adjacent to R301 at the bottom of TM4. Mutations in this region of TM4 have profound effects on pLGIC desensitization (48, 49, 52, 53). R117 and R123 are located at the extracellular end of TM4, and boundary POPG with headgroups that interact with these residues have acyl chains that make contacts with intrasubunit sites on both sides of TM4 (Fig. 8B and 5B). Sites equivalent to R117 and R123 in GLIC were previously found to be occupied by a phospholipid and docosahexaenoic acid (DHA), respectively (24, 25). The polyunsaturated fatty acid, DHA, was found to increase desensitization in GLIC (25). Therefore, it is possible that the exact lipid structure occupying these sites results in different effects. Our results raise the hypothesis that lipids with polyunsaturated acyl chains or certain sterols (54) exert the opposite effect of activating phospholipids by acting as competitive antagonists.

In summary, the anionic phospholipid, POPG, decreases desensitization in the pLGIC, ELIC. POPG specifically binds to and stabilizes ELIC by interacting with interfacial arginine residues. Our results strongly suggest that binding of POPG at specific sites modulates receptor desensitization.

## Materials and Methods

### Mutagenesis, expression and purification of ELIC

pET26-MBP-ELIC was a gift from Raimund Dutzler (Addgene plasmid # 39239) and was used for WT ELIC expression and generation of mutants. Site-directed mutagenesis was performed by the standard Quikchange approach, and confirmed by Sanger sequencing (Genewiz, Plainfield, NJ). WT and mutant ELIC was expressed as previously described (25, 55) in OverExpress^TM^ C43 (DE3) *E. coli* (Lucigen, Middleton, WI). Cultures were grown in Terrific Broth (Sigma, St. Louis, MO) and induced with 0.1 mM IPTG for ∼16 hours at 18 °C. Pelleted cells were resuspended in Buffer A (20 mM Tris pH 7.5, 100 mM NaCl) with cOmplete EDTA-free protease inhibitor (Roche, Indianapolis, IN), and lysed using an Avestin C5 emulsifier at ∼15,000 psi. Membranes were collected by ultracentrifugation, resuspended in Buffer A, solubilized in 1 % DDM (Anatrace, Maumee, OH), and incubated with amylose resin (New England Biolabs, Ipswich, MA) for 2 hours. The resin was washed with 20 bed volumes of Buffer A, 0.02% DDM, 0.5 mM tris(2-carboxyethyl)phosphine (TCEP) and 1 mM EDTA, and eluted with Buffer A, 0.02% DDM, 0.05 mM TCEP, and 40 mM maltose. Eluted protein was digested overnight with HRV-3C protease (Thermo Fisher, Waltham, MA) (10 units per mg ELIC) at 4 °C, and injected on a Sephadex 200 10/300 (GE Healthcare Life Sciences, Pittsburgh, PA) size exclusion column in Buffer A, 0.02% DDM.

### Native MS measurements

Native MS analysis was similar to previous descriptions for other membrane proteins (27). For analysis of ELIC in DDM, 30 μl of purified protein in 0.02% DDM at ∼1 mg/ml was buffer exchanged into 200 mM ammonium acetate pH 7.5 and 0.02% DDM using Biospin 6 gel filtration spin columns (Bio-Rad, Hercules, CA). 2 μl of buffer exchanged ELIC was loaded into a borosilicate capillary emitter (Thermo Scientific, Waltham, MA), and analyzed by static nanospray on a Thermo QExactive EMR mass spectrometer. The following parameters were used to resolve the ELIC pentamer and minimize dissociation into tetramer and monomer: capillary voltage of 1.2 kV, capillary temperature of 200 °C, ion transfer optics set with the injection flatapole, inter-flatapole lens, bent flatapole, transfer multiple as 8, 7, 6, 4 V, respectively, resolution 8,750, AGC target 3 x 10^6^, trap pressure set to maximum, CID 200 V, and CE 100 V. For analysis of ELIC in C10E5, ELIC delipidated by injecting 300 μg onto a Sephadex 200 10/300 column (GE Healthcare) at 0.5 ml/min pre-equilibrated with Buffer A, 10% glycerol, and 0.06% C10E5 (Anatrace). 30 μl aliquots were then buffer exchanged to 100 mM ammonium acetate pH 7.5, 0.06% C10E5 using Biospin 6 columns, and diluted to 0.2 mg/ml. MS measurements on the QExactive EMR were performed with the parameters listed above except: capillary temperature 100 °C, CID 75 V and CE 200 V. For lipid binding measurements, stocks of POPG lipid were prepared at 2x the concentration of POPG being tested in 100 mM ammonium acetate pH 7.5 and 0.06% C10E5. Lipid stocks were then mixed with 0.4 mg/ml ELIC in a 1:1 volume ratio for a final concentration of 1 μM ELIC, and samples were analyzed after >5 min incubation.

MS spectra were deconvoluted using UniDec (56); deconvolution of spectra with bound lipid was restricted to the +26 to +22 charge states (Supplementary Fig. 3). Peak heights of apo and lipid-bound species were extracted from UniDec, and analyzed by two approaches. The average number of bound lipids was determined by the following relationship:

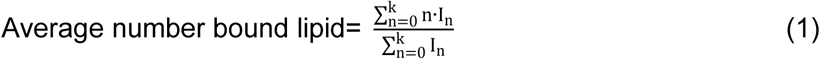

where n is the number of bound lipids and I_n_ is the deconvoluted peak height of ELIC with n bound lipids. Peak heights of apo and lipid-bound species were also plotted as mole fraction versus the number of bound lipids (Supplementary Fig. 4). These data were fit with a binomial binding model, which assumes that there are N sites each with equal affinity, K. The probability, p, that a site is occupied at the concentration of a given lipid, A, is defined as:

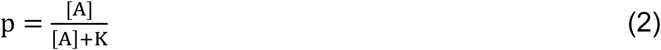

Then, the probability (B) that q sites are occupied out of N total sites is given by the binomial probability function:

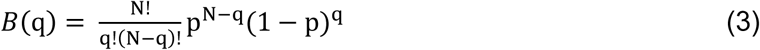

B(q) was used to determine the mole fraction of each lipid-bound species at a given [A], which was used to fit the native MS data in Excel across all [A] by setting K constant and varying N or vice versa.

### Thermal stability assay

Purified WT ELIC in C10E5 (Buffer A, 0.06% C10E5) was diluted to 1 μM in the absence or presence of various concentrations of phospholipid. Samples were analyzed without and with heating in the absence or presence of phospholipid. Analysis of protein thermal stability was performed by injecting 90 μl of sample on a size exclusion column (Sephadex 200 10/300), and measuring the amplitude of the pentamer peak as previously described (57, 58). Heating was performed for 15 min at 32 °C, which resulted in a ∼85% decrease in the pentamer amplitude compared to 4°C. The stabilizing effect of a phospholipid was quantified as the pentamer amplitude in the presence of phospholipid (heated) divided by control (heated).

### Excised patch-clamp recordings from giant liposomes

ELIC WT and mutants were reconstituted into giant liposomes as previously described with some modifications (59). Three liposome preparations were used in this study: 1) 25% POPG (consists of 50% POPC/ 25% POPE/ 25% POPG), 2) 12% POPG (consists of 60% POPC/ 28% POPE/12% POPG), and 3) 40% POPG (consists of 35% POPC/ 25% POPE/ 40% POPG). These liposome compositions were chosen to vary POPG mole% while optimizing lipid mixtures to obtain ideal giant liposomes for patch-clamping. This was achieved by varying POPG mole% and POPC mole% inversely. Condition #1 was used for WT and all mutants, and conditions #2 and #3 were used in WT. Liposomes were prepared by drying 15-20 mg of lipid mixtures in chloroform using N_2_ in a round bottom flask and then overnight in a vacuum dessicator. Dried lipids were rehydrated at 5 mg/ml in 10 mM MOPS pH 7, and 150 mM NaCl (MOPS buffer), subjected to ten freeze-thaw cycles ten, and then small unilamellar liposomes were formed by extrusion using a 400 nm filter (Avanti Lipids, Alabaster, AL) and bath sonication (30 sec x 5). 5 mg of liposomes in 1 ml were destabilized by adding DDM to 0.2% and rotating for 1 hour at room temperature followed by 0.3-0.5 mg of ELIC WT or mutants at ∼4-5 mg/ml and incubation for 30 min. To remove DDM, SM-2 Bio-beads (Bio-Rad) were added in five batches (30, 30, 50, 100, and 100 mg). The first three batches were added each hour along with 1 ml of MOPS buffer to make a final volume of 4 ml while rotating at room temperature. After adding the first 100 mg batch, the proteoliposomes were rotated overnight at 4 °C, followed by the last 100 mg the next day for 3 hours at room temperature. Proteoliposomes were harvested by ultracentrifugation at 150,000 x *g* for 1 hour at 4 °C, and the pellet resuspended with 80 μl of MOPS buffer for a lipid concentration of ∼50 mg/ml. Giant liposomes were formed by drying 10 μl of proteoliposomes on a glass coverslip in a desiccator for 3-5 hours at 4 °C followed by rehydration with 60 μl of MOPS buffer overnight at 4 °C and at least 2 hours at room temperature the next day. Giant liposomes were resuspended by pipetting and then applied to a petri dish with MOPS buffer.

Patch-clamp recordings were performed using borosilicate glass pipettes pulled to ∼2-3 MΩ using a P-2000 puller (Sutter instruments, Novato, CA). Pipettes were filled with 10 mM MOPS pH 7, 150 mM NaCl, and 0.5 mM BaCl_2_. Excised patches (the orientation of ELIC in the liposomes is not known; therefore, these patches are not defined as outside-out or inside-out) were held at −60 mV, and bath solutions consisted of 10 mM MOPS pH 7, 150 mM NaCl, 0.5 mM BaCl_2_, 1 mM dithiothreitol (DTT), and varying concentrations of cysteamine. DTT was added to the bath solution to prevent cysteamine oxidation. Rapid solution exchange was achieved with a three-barreled flowpipe mounted and adjusted by to a SF-77B fast perfusion system (Warner Instrument Corporation, Hamden, CT). Liquid junction current at the open pipette tip demonstrated 10-90% exchange times of <10 ms. Data was collected at 20 kHz using an Axopatch 200B amplifier (Molecular Devices, San Jose, CA) and a Digidata 1322A (Molecular Devices) with Axopatch software, and a low pass Bessel filter of 10 kHz was applied to the currents. Analysis of currents was performed with Clampfit 10.4.2 (Molecular Devices). Activation currents were fit to a single exponential equation, and desensitization currents were fit to both single and double exponential equations. The majority of desensitization currents were best fit with a double exponential, and weighted time constants were derived using the following calculation:

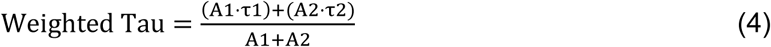

where A1 and A2 are the weighted coefficients of the first and second exponential components. The reported weighted average time constants are averages of weighted time constants from double exponential fits and time constants from single exponential fits. Peak cysteamine dose response curves were fit to a Hill equation, keeping n constant at 2, which provided a reasonable fit for all data sets. All statistical comparisons were made using a one-way ANOVA with post-hoc Tukey HSD test.

### Stopped-flow fluorescence recordings

The fluorescence-based sequential-mixing stopped-flow assay was carried out with an SX20 stopped-flow spectrofluorimeter (Applied Photophysics, Leatherhead, UK) at 25 °C. To reconstitute ELIC into large unilamellar vesicles (LUVs), 15 mg of lipids (POPC or POPC:POPE:POPG 2:1:1) were dried in glass vials to a thin film under a constant N_2_ stream. Lipids were further dried under vacuum overnight. The next day, lipids were rehydrated in reconstitution buffer (1114 µl of 15 mM Hepes, 150 mM NaNO_3_, pH 7). 33 mg CHAPS were added stepwise while sonicating lipids in a bath sonicator until the solution was clear. 1057 µl of a 75 mM ANTS stock solution (in ddH_2_O, pH 7) was added together with purified ELIC (1 µg/mg lipid), mixed and incubated for 20 min. Detergent removal was initiated by addition of 0.7 g SM-2 BioBeads (BioRad) in assay buffer (10 mM Hepes, 140 mM NaNO_3_, pH 7). The reconstitution mix was incubated for 2.5 h at 21 °C under gentle agitation. The liposome-containing supernatant was transferred to a new glass tube and stored overnight at 13 °C. The liposome solution was sonicated in a bath sonicator for 30 s and extruded through a 0.1 µm membrane (Whatman) using a mini-extruder (Avanti Polar lipids). Extra-vesicular ANTS were removed with a 10 ml desalting column (PD-10, GE Lifesciences). Right before the assay, liposomes were diluted 5-fold in assay buffer to ensure a good signal to noise ratio.

For the assay, ELIC-containing liposomes were mixed 1:1 with pre-mix buffer (assay buffer supplemented with 10 mM cysteamine to reach 5 mM after mixing) and incubated for defined amounts of time (10 ms to 25 s). A second 1:1 mixing step was performed with quenching buffer (10 mM Hepes, 90 mM NaNO_3_, 50 mM TlNO_3_, pH 7). ANTS fluorescence was excited at 360 nm and the integral fluorescence above 420 nm was recorded for 1 s. For each delay time, at least 8 repeats under identical conditions were performed.

To analyze the data, each repeat was visually inspected and outliers were removed. Each remaining repeat was then fitted to a stretched exponential (Equation 5) and the rate of Tl+ influx was determined at 2 ms (Equation 6).

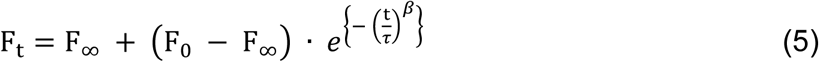

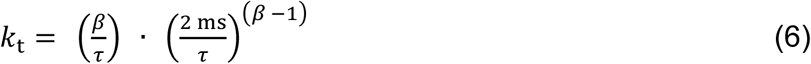

with F_t_, F_∞_, F_0_ being the fluorescence at time t, the final fluorescence and the initial fluorescence, respectively. t is the time (in s), τ the time constant (in s) and *β* the stretched exponential factor. *k*_t_ is the calculated rate (in s^-1^) of Thallium influx at 2 ms.

The rate constants were averaged and the mean and standard deviations were determined and plotted (Fig 2F). The experiments were repeated for each lipid composition using three independent reconstitutions. The rates and standard deviations were averaged and plotted as function of the delay time. The time course was fitted according to a double-exponential function to obtain the rates of activation and desensitization.

### Lipid extraction and MS analysis

Lipids were extracted using a Bligh-Dyer extraction (60). Briefly, 100 μg of purified ELIC in DDM and 150 μg of *E. coli* membranes derived from cell cultures transformed and induced for ELIC expression, respectively, were mixed with 1 ml chloroform, 2 ml methanol and 0.8 ml water, and vortexed for 1 min, followed by an additional 1 ml chloroform and 1 ml water, and vortex for 3 min. The samples were centrifuged for 3 min at 500 x *g*, and the lower organic phase removed for analysis, using a Thermo Scientific LTQ Orbitrap Velos mass spectrometer. Lipid extracts were loop injected (1.5 μl/min) using a syringe pump that delivered a continuous flow of methanol at 15 μl/min into the ESI source. High resolution (R = 100,000 at m/z 400) MS and MS/MS analyses were performed in negative ion mode. The skimmer of the ESI source was set at ground potential, electrospray voltage 4 kV, capillary temperature 300 °C, AGC target 5 x 10^4^, and maximum injection time 50 ms. MS^n^ experiments for identification of lipid structures were carried out with an optimized relative collision energy of 32%, activation q value of 0.25, activation time of 10 ms, and mass selection window of 1 Da.

### Coarse-grained simulations of ELIC

All simulations reported here used the MARTINI 2.2 (61) coarse-grained topology and force field. The crystal structure of ELIC (PDB 3RQW) (62) was coarse-grained using MARTINI martinize.py script. Secondary structural restraints were constructed using martinize.py while imposed through Gromacs (63). Conformational restraints were preserved through harmonic bonds between backbone beads less than 0.5 nm apart with a coefficient of 900 kJ mol^-1^. Pairs were determined using the ElNeDyn algorithm (64). Membranes were constructed using the MARTINI script insane.py (61). The insane.py script randomly places lipids throughout both inner and outer membranes and embeds selected proteins into the membrane. Two series of simulations were developed, the first using POPE and POPG, and the second POPC and POPG. Box sizes were about 30 x 30 x 25 nm^3^ and each simulation box contained about 3000 lipids.

Molecular dynamics simulations were carried out using GROMACS 5.1.4 (63). All systems were run using van der Waals (vdW) and electrostatics in cutoff and reaction-field, respectively, with a dielectric constant of *ε* = 15. vdW and electrostatics used a cutoff length of 1.1 nm as defined in current MARTINI build specifications. Energy minimizations were performed for about 30,000 steps. All systems were run for short equilibration steps. Canonical ensembles (NVT) were run for 100 ps using Berendsen thermostat set to 323 K with the temperature coupling constant set to 1 ps. Isothermal-Isobaric ensemble (NPT) equilibration was run for 5000 ps using Berendsen thermostat and barostat. The thermostat was set to 323 K with the temperature coupling constant set to 1 ps, and the barostat was set to a pressure coupling constant of 3 ps with a compressibility of 3 x 10^-5^ bar^-1^ holding at 1 bar. Molecular dynamics were carried out using NPT ensemble and were simulated for 15 μs with a time step of 0.015 ps using v-rescale thermostat set to 323 K and a temperature coupling constant of 1 ps. Membranes consisting of POPE used the Parrinello-Rahman barostat, and membranes consisting of POPC used the Berendsen barostat, both under semi-isotropic coupling. The reference pressure was set to 1 bar, the compressibility 3×10^-4^ bar^-1^, and the pressure coupling constant 1 ps.

Annular lipids were determined using the annular lipid metric B:

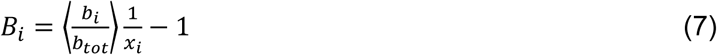

where *b_i_* is the instantaneous number of boundary lipids of species *i*, *b_tot_* is the instantaneous total number of boundary lipids, *x*_*i*_ is the overall (bulk) fraction of species *i* and the brackets represent an average over time and replicas. *B_i_* < 0 and *B_i_* > 0 indicate enrichment and depletion of species *i*, respectively, relative to the abundance in the bulk membrane. A given lipid was counted as a boundary lipid if it was within 6 Å of the ELIC transmembrane domain.

Two dimensional lipid density distributions around a central ELIC pentamer were calculated for each leaflet using polar coordinates (28). For every sampled frame, all lipids of species *i* were separated into leaflets. For all *i* lipids in a given leaflet, the vector separating the phosphate beads from ELIC center was calculated and projected onto the membrane plane. The two-dimensional separation vector was then used to assign the lipid to the appropriate polar bin of radial bin width 4Å and angular bin width 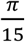. The area density in each bin was averaged over time and replicas.

## Acknowledgements

We are grateful to Alex Evers, Joe Henry Steinbach, Christopher Lingle and Gustav Akk for helpful discussions and edits regarding this study and the preparation of the manuscript. We also acknowledge Arthur Laganowsky and Yang Liu for guidance with regard to sample preparation of ELIC for native MS measurements. We are indebted to Michael Gross at the Washington University NIH/NIGMS-supported biomedical mass spectrometry resource for use of the Thermo QExactive EMR mass spectrometer, and Christopher Lingle for use of a patch-clamp rig for electrophysiology recordings. Computational resources were provided through the Rutgers Discovery Informatics Institute.

## Supplementary Data

**Supplementary Fig 1.**
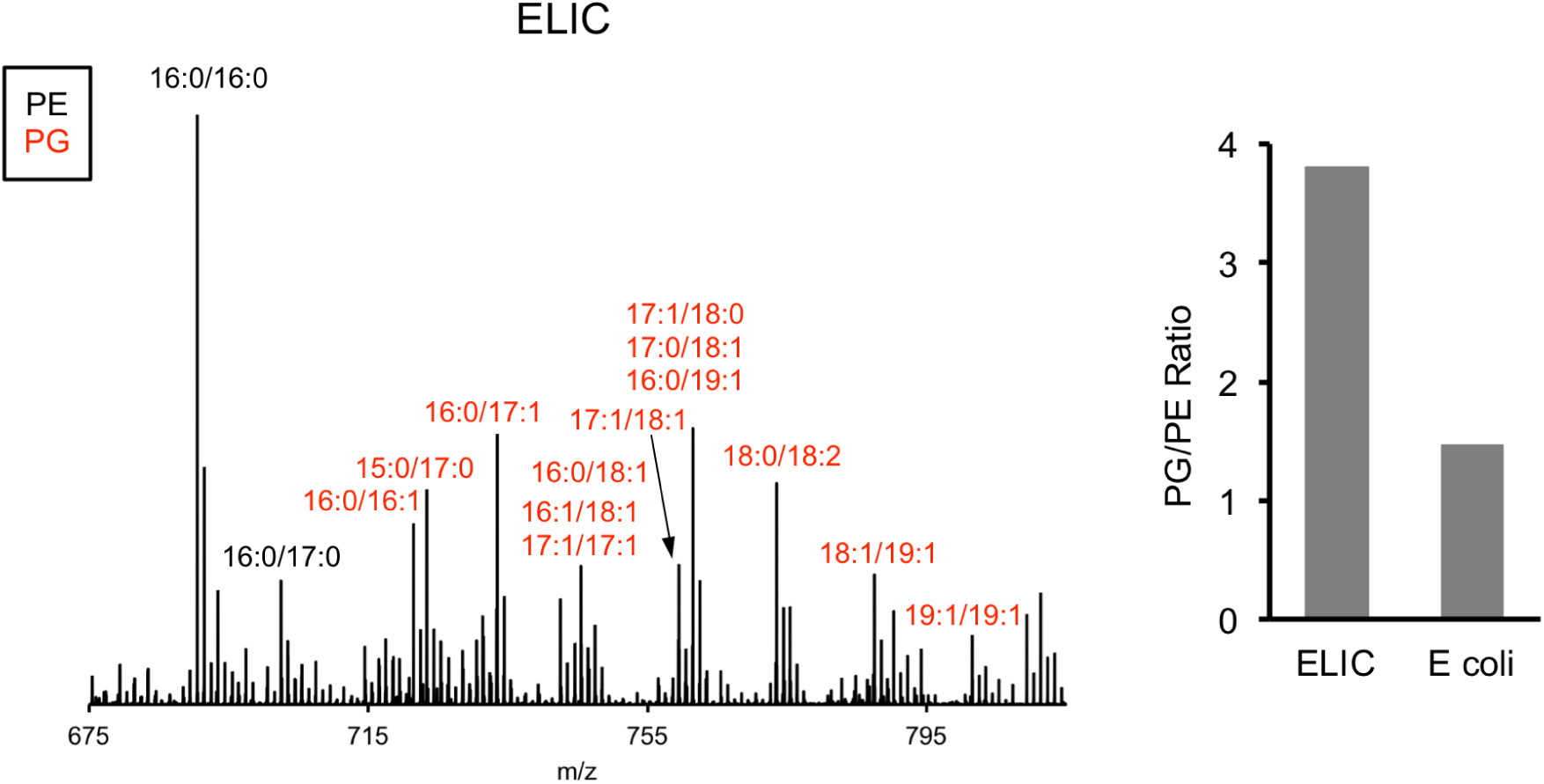
MS1 spectra of lipid extract from purified ELIC in DDM and *E. coli* membranes. Labeled peaks correspond to PG (red) and PE (black) phospholipids with specific acyl chain combinations determined from MS2 fragmentation. *Right:* Graph shows quantification of the intensity of all PG species relative to PE species for ELIC and *E. coli* membrane samples.

**Supplementary Fig. 2.**
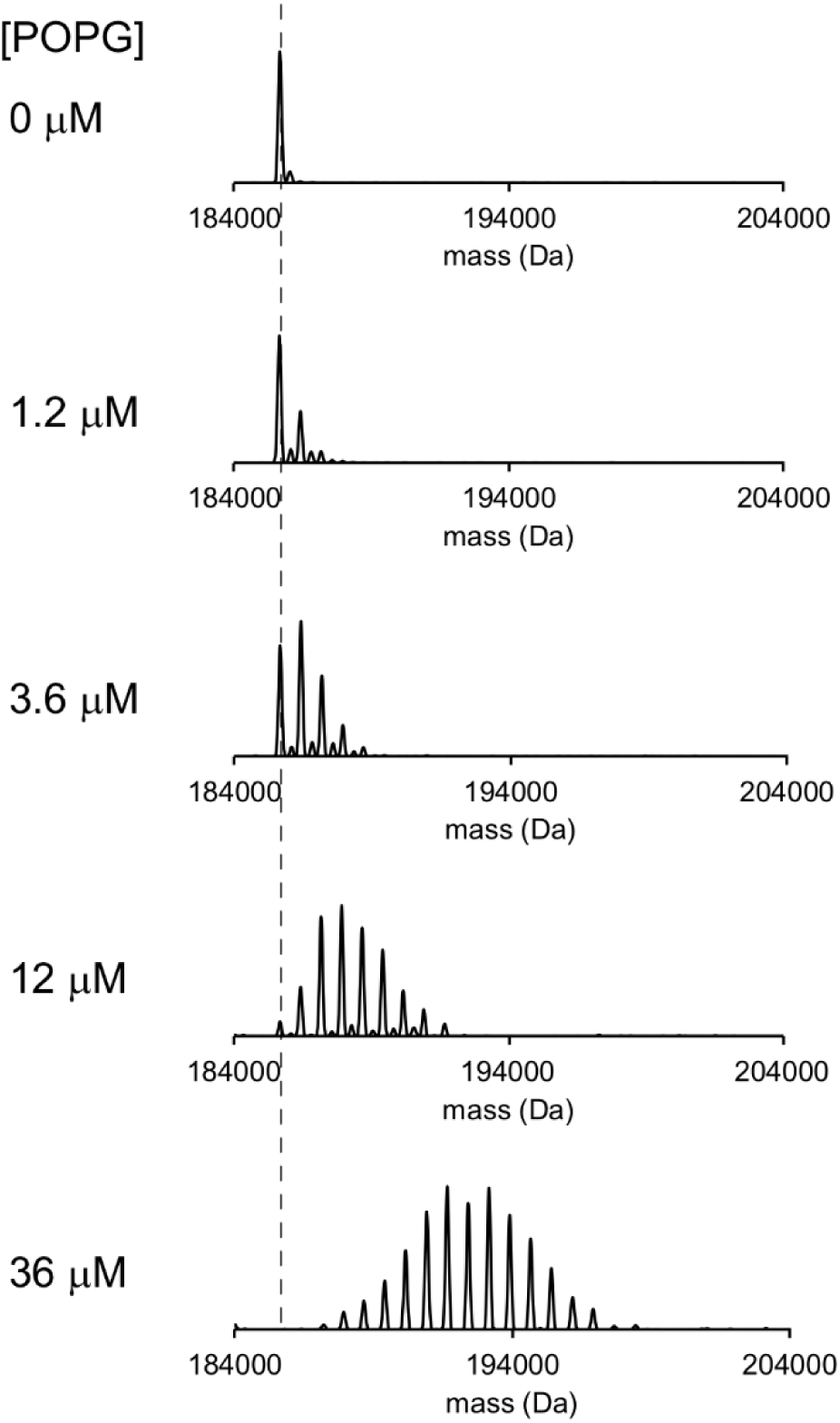
Representative deconvoluted spectra of 1 μM ELIC in C10E5 with increasing concentration of POPG. Dashed line indicates mass of apo ELIC.

**Supplementary Fig. 3.**
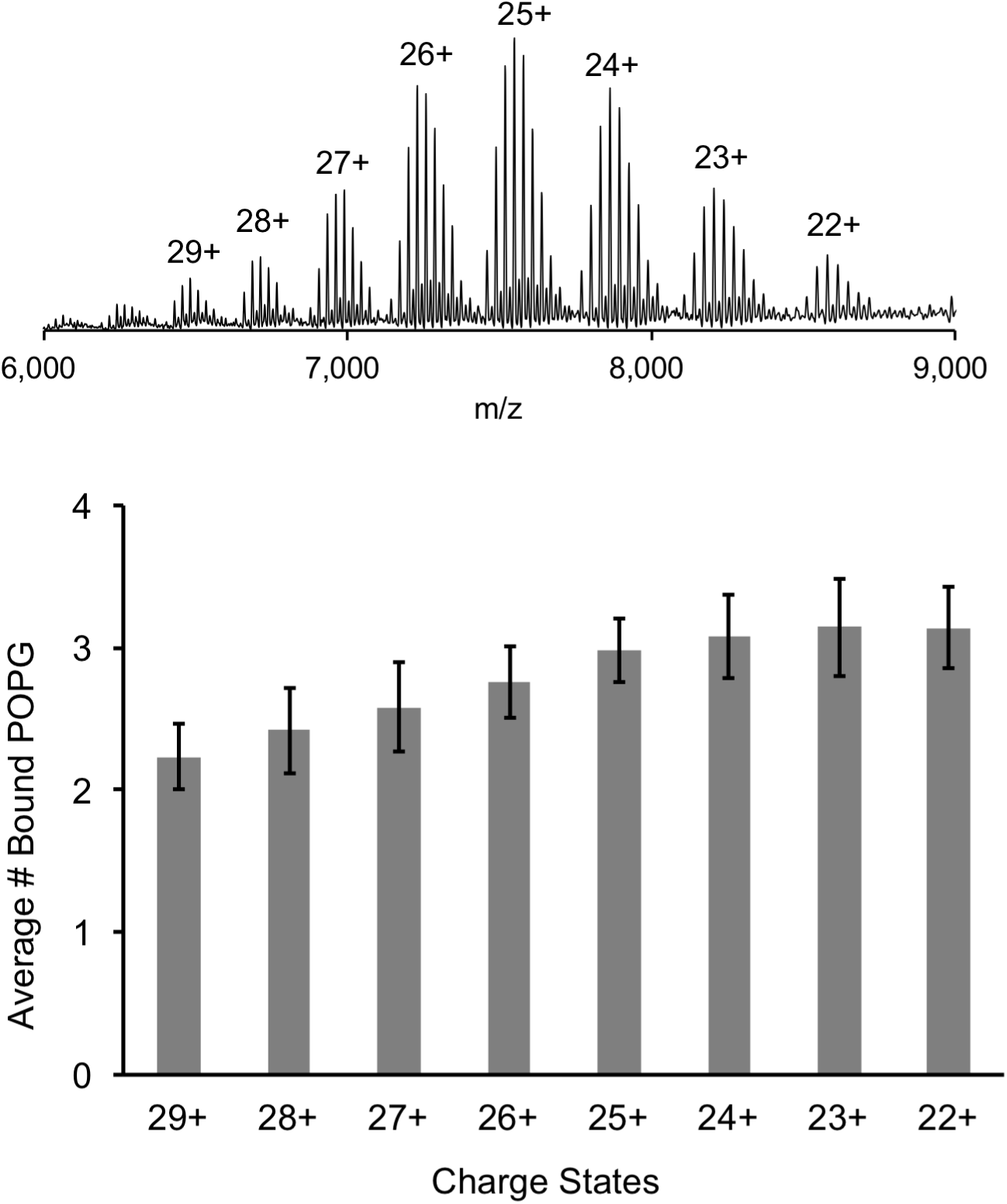
Comparison of lipid binding at different charge states. *Top:* Representative full native spectrum of the ELIC pentamer with 12 μM POPG with each charge state labeled. *Bottom:* Quantification of the average number of bound POPG to the ELIC pentamer at each charge state (n=13, ±SD).

**Supplementary Fig. 4.**
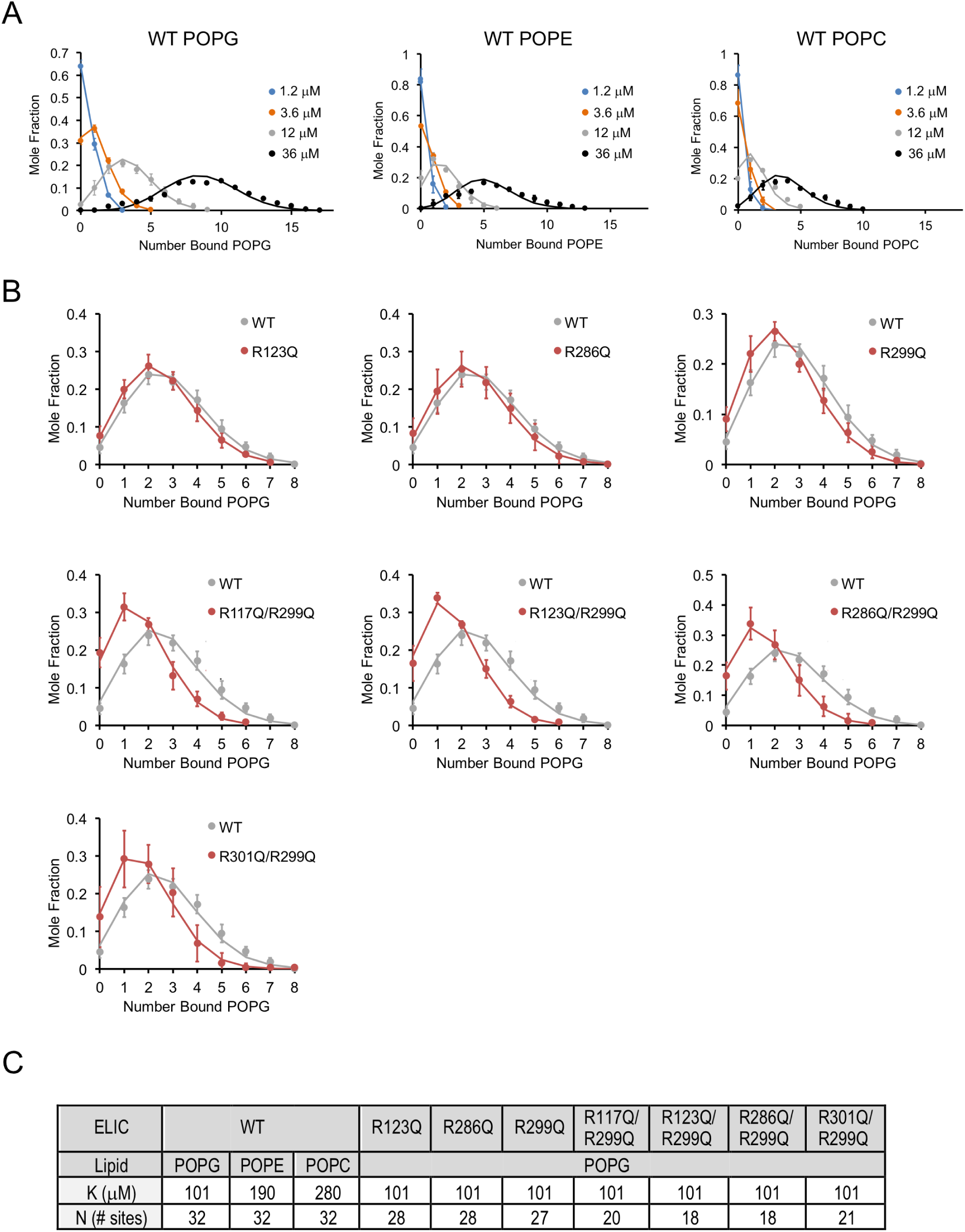
Lipid binding data fit to binomial distributions. (**A**) Plots of mole fraction of phospholipid-bound ELIC derived from native MS experiments with varying concentrations of phospholipid (circles, n=3-6, ±SD). Solid lines show global fits from a binding model based on a binomial distribution with 32 sites (N) of equal affinity (K) using K as shown in (C). (**B**) Plots of mole fraction of POPG-bound ELIC WT and mutants at 12 μM POPG (circles, n=3-6, ±SD). Solid lines show fits as in (A) in which K is held constant at 102 μM and N is varied as indicated in (C). (**C**) Table showing dissociation constants (K) and number of sites (N) used in fits shown in (A) and (B).

**Supplementary Fig. 5.**
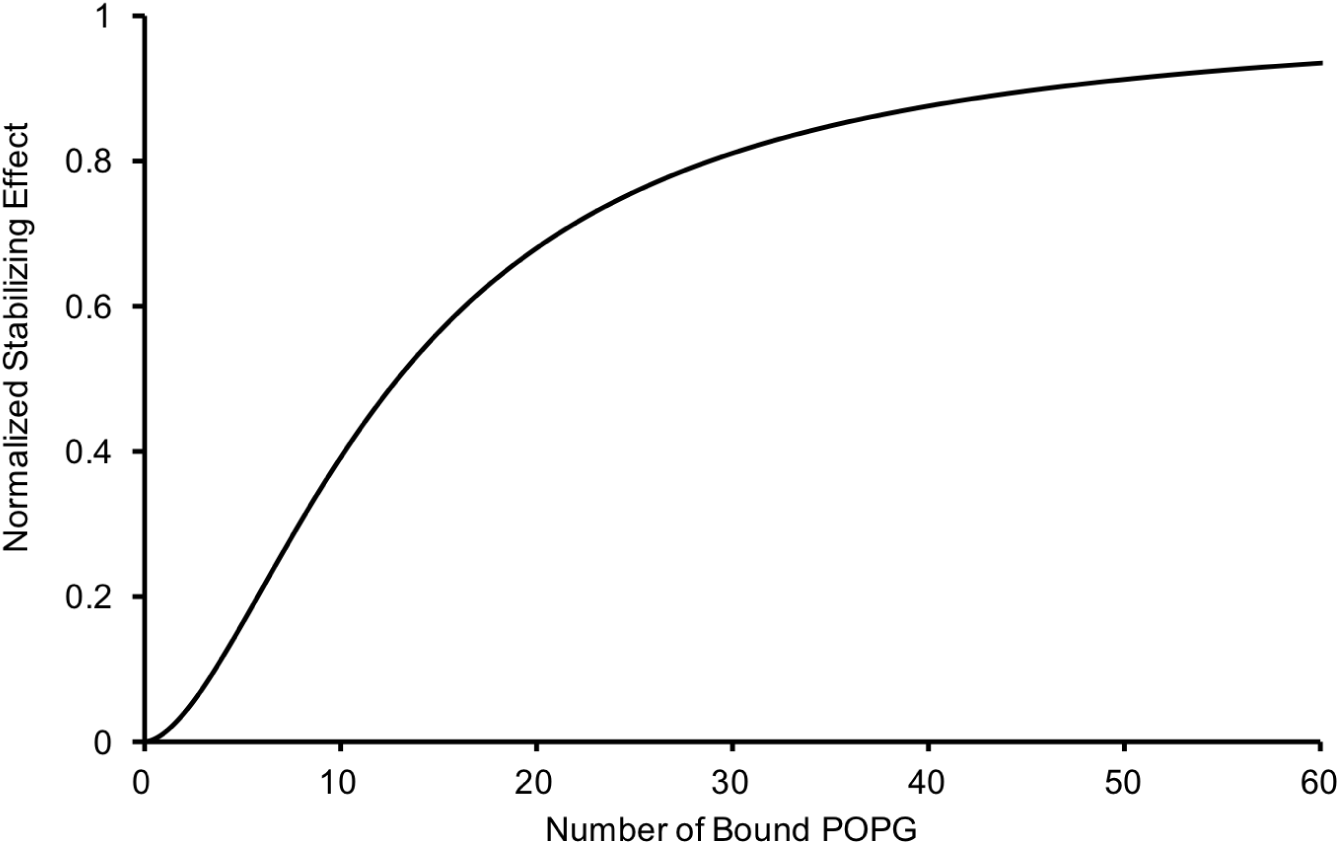
Relationship of the thermal stabilizing effect of POPG vs the average number of bound POPG derived from equating concentration of POPG from the sigmoid functions used to fit the POPG binding (Fig. 1B) and thermal stability data (Fig. 2A). The resulting relationship is: 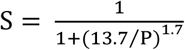, where P is the average number of bound POPG and S is the normalized thermal stabilizing effect.

**Supplementary Fig. 6.**
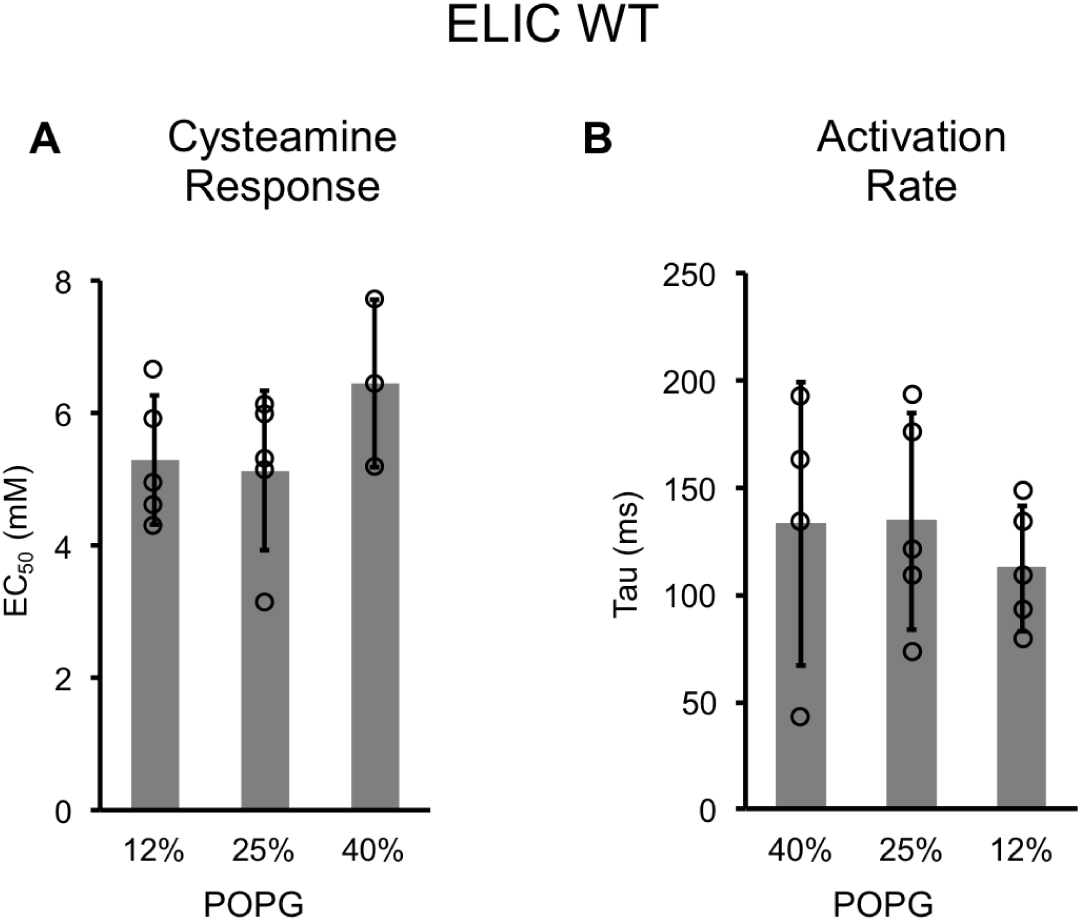
Channel properties of WT ELIC responses to cysteamine. (**A**) EC_50_ for peak responses to cysteamine of WT ELIC in giant liposomes of varying mole% POPG (n=3-5, ±SD). (**B**) Activation time constants (τ) derived from single exponential fits of WT ELIC in response to 30 mM cysteamine in giant liposomes of varying mole% POPG (n=4-5, ±SD).

**Supplementary Fig. 7.**
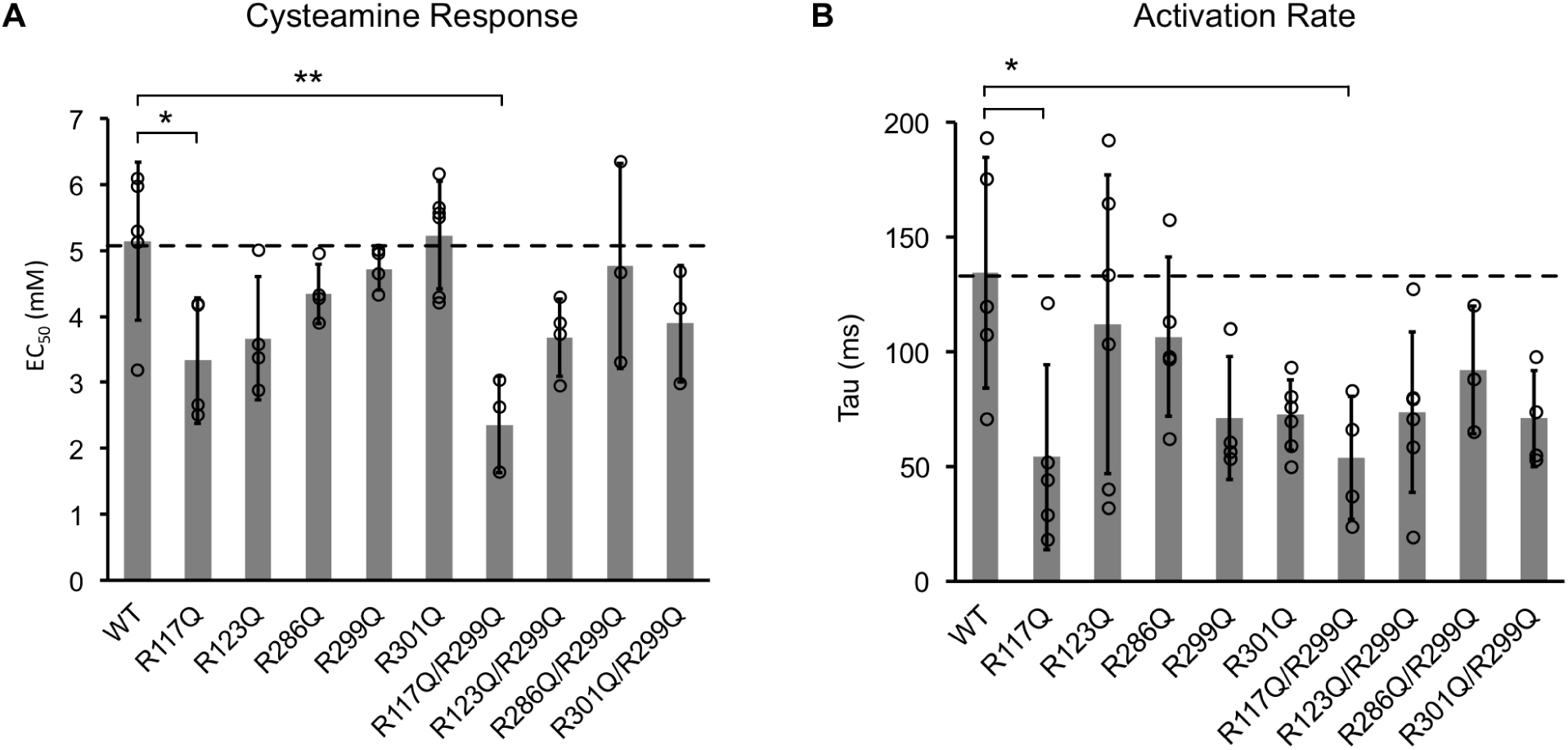
Channel properties of ELIC WT and mutant responses to cysteamine in giant liposomes of 25 mole% POPG. (**A**) Graph of EC_50_ for peak responses to cysteamine of ELIC WT and mutants (n=4-7, ±SD, *p<0.05, **p<0.01). (**B**) Activation time constants (tau) of ELIC WT and mutants in response to 30 mM cysteamine (n=4-7, ±SD, *p<0.05).

**Supplementary Table 1:**
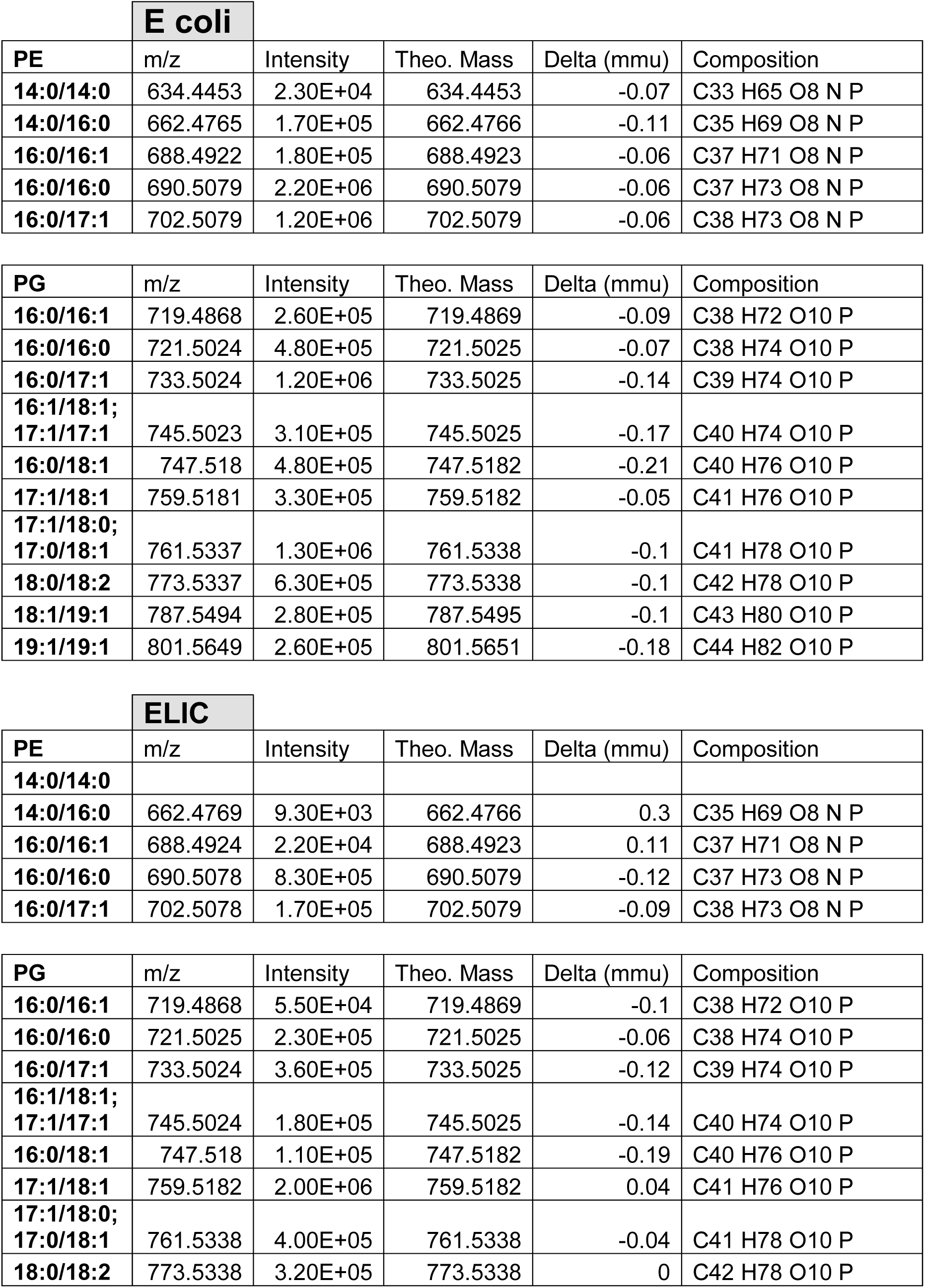
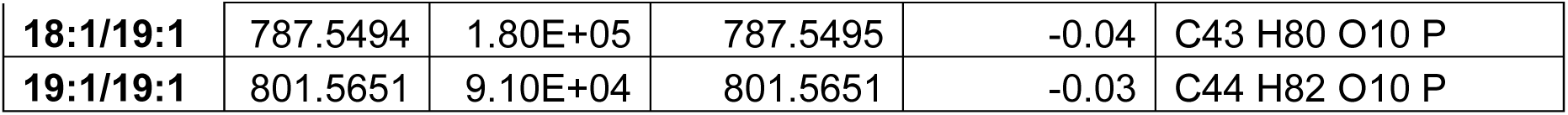
Phophatidylethanolamine and phosphatidylglycerol species identified in lipid extracts by MS/MS. Table shows m/z, intensity, mass and mass accuracy of each phospholipid species.

**E coli**
| PE | m/z | Intensity | Theo. Mass | Delta (mmu) | Composition |
| --- | --- | --- | --- | --- | --- |
| 14:0/14:0 | 634.4453 | 2.30E+04 | 634.4453 | -0.07 | C33 H65 O8 N P |
| 14:0/16:0 | 662.4765 | 1.70E+05 | 662.4766 | -0.11 | C35 H69 O8 N P |
| 16:0/16:1 | 688.4922 | 1.80E+05 | 688.4923 | -0.06 | C37 H71 O8 N P |
| 16:0/16:0 | 690.5079 | 2.20E+06 | 690.5079 | -0.06 | C37 H73 O8 N P |
| 16:0/17:1 | 702.5079 | 1.20E+06 | 702.5079 | -0.06 | C38 H73 O8 N P |

| PG | m/z | Intensity | Theo. Mass | Delta (mmu) | Composition |
| --- | --- | --- | --- | --- | --- |
| 16:0/16:1 | 719.4868 | 2.60E+05 | 719.4869 | -0.09 | C38 H72 O10 P |
| 16:0/16:0 | 721.5024 | 4.80E+05 | 721.5025 | -0.07 | C38 H74 O10 P |
| 16:0/17:1 | 733.5024 | 1.20E+06 | 733.5025 | -0.14 | C39 H74 O10 P |
| 16:1/18:1;<br>17:1/17:1 | 745.5023 | 3.10E+05 | 745.5025 | -0.17 | C40 H74 O10 P |
| 16:0/18:1 | 747.518 | 4.80E+05 | 747.5182 | -0.21 | C40 H76 O10 P |
| 17:1/18:1 | 759.5181 | 3.30E+05 | 759.5182 | -0.05 | C41 H76 O10 P |
| 17:1/18:0;<br>17:0/18:1 | 761.5337 | 1.30E+06 | 761.5338 | -0.1 | C41 H78 O10 P |
| 18:0/18:2 | 773.5337 | 6.30E+05 | 773.5338 | -0.1 | C42 H78 O10 P |
| 18:1/19:1 | 787.5494 | 2.80E+05 | 787.5495 | -0.1 | C43 H80 O10 P |
| 19:1/19:1 | 801.5649 | 2.60E+05 | 801.5651 | -0.18 | C44 H82 O10 P |

**ELIC**
| PE | m/z | Intensity | Theo. Mass | Delta (mmu) | Composition |
| --- | --- | --- | --- | --- | --- |
| 14:0/14:0 |  |  |  |  |  |
| 14:0/16:0 | 662.4769 | 9.30E+03 | 662.4766 | 0.3 | C35 H69 O8 N P |
| 16:0/16:1 | 688.4924 | 2.20E+04 | 688.4923 | 0.11 | C37 H71 O8 N P |
| 16:0/16:0 | 690.5078 | 8.30E+05 | 690.5079 | -0.12 | C37 H73 O8 N P |
| 16:0/17:1 | 702.5078 | 1.70E+05 | 702.5079 | -0.09 | C38 H73 O8 N P |

**Supplementary Table 1:** Phosphatidylethanolamine and phosphatidylglycerol species identified in lipid extracts by MS/MS. Table shows m/z, intensity, mass and mass accuracy of each phospholipid species.
| PG | m/z | Intensity | Theo. Mass | Delta (mmu) | Composition |
| --- | --- | --- | --- | --- | --- |
| 16:0/16:1 | 719.4868 | 5.50E+04 | 719.4869 | -0.1 | C38 H72 O10 P |
| 16:0/16:0 | 721.5025 | 2.30E+05 | 721.5025 | -0.06 | C38 H74 O10 P |
| 16:0/17:1 | 733.5024 | 3.60E+05 | 733.5025 | -0.12 | C39 H74 O10 P |
| 16:1/18:1;<br>17:1/17:1 | 745.5024 | 1.80E+05 | 745.5025 | -0.14 | C40 H74 O10 P |
| 16:0/18:1 | 747.518 | 1.10E+05 | 747.5182 | -0.19 | C40 H76 O10 P |
| 17:1/18:1 | 759.5182 | 2.00E+06 | 759.5182 | 0.04 | C41 H76 O10 P |
| 17:1/18:0;<br>17:0/18:1 | 761.5338 | 4.00E+05 | 761.5338 | -0.04 | C41 H78 O10 P |
| 18:0/18:2 | 773.5338 | 3.20E+05 | 773.5338 | 0 | C42 H78 O10 P |
| **18:1/19:1** | 787.5494 | 1.80E+05 | 787.5495 | -0.04 | C43 H80 O10 P |
| **19:1/19:1** | 801.5651 | 9.10E+04 | 801.5651 | -0.03 | C44 H82 O10 P |

